# Systems Analysis of Carboxylate Transport and Oxidation Pathways in Cardiac Mitochondria

**DOI:** 10.64898/2026.02.25.708012

**Authors:** Nicole L. Collins, Santosh Dasika, Françoise Van den Bergh, Jason Bazil, Daniel A. Beard

## Abstract

Experimental assessment and computational modeling were used to analyze substrate transport, tricarboxylic acid cycle kinetics, and oxidative phosphorylation in suspensions of purified cardiac mitochondria. The kinetics of ATP synthesis and carbohydrate oxidation, including during hypoxia and reoxygenation, were investigated using various substrate combinations and conditions. Model simulations fit to transient respiration and NAD(P)H measurements reveal novel insights into pyruvate dehydrogenase regulation, regulation of mitochondrial leak, and the clearance of oxaloacetate during respiration on succinate. High concentrations of succinate induced increased mitochondrial leak respiration driven in part by ROS-activated uncoupling. Oxidative phosphorylation under succinate-fueled respiration was inhibited by rapid buildup of oxaloacetate, inhibiting succinate dehydrogenase. Malic enzyme and oxaloacetate decarboxylase activities represent potenital pathways for removal of oxaloacetate, with glutamate further enhancing clearance. The developed model captures the observed transient behaviors as well as steady-state relationships between ATP synthesis rate and phosphate metabolite levels, lending a new systems-level understanding of mitochondrial energy metabolism. In sum, these findings offer a framework for simulating and interpreting mitochondrial function in vitro and in vivo.

**Key Points:** This study uses experiments and computer simulations to probe the interactions between substrate transport processes, TCA cycle kinetics, redox state, and oxidative ATP synthesis in cardiac mitochondria.

The developed kinetic model simulates mitochondrial metabolism in vitro and represents a framework for integrative modeling of cardiac energy metabolism.

Model-based analysis identifies a kinetic model of pyruvate dehydrogenase (PDH) deactivation during leak-state respiration and activation during oxidative phosphorylation.

High levels of cation leak during respiration on succinate are explained by a ROS-dependent activation of uncoupling.

## Introduction

Mitochondria are at the center of energy metabolism and contribute numerous essential biochemical pathways. The final steps in the oxidative breakdown of carbohydrates, fatty acids, and amino acids occur in the tricarboxylic acid (TCA) cycle, via which reducing equivalents are generated to fuel oxidative ATP synthesis. The TCA cycle also supplies key intermediates that serve as precursors for synthesis of fatty acids and amino acids [1, 2]. Furthermore, certain TCA cycle intermediates exported from the mitochondrial matrix play roles as extramitochondrial signaling molecules and as substrates for epigenetic modification [2]. While the net canonical reaction of the TCA cycle consists of complete oxidation of acetate (as acetyl-coenzyme A) to carbon dioxide and coenzyme A, several intermediates enter and exit the TCA cycle through metabolic pathways shared between the mitochondrial matrix and the cytosol. TCA cycle intermediates and related precursors imported and exported from the mitochondrial matrix include pyruvate, citrate, a-ketoglutarate, succinate, malate, glutamate, and aspartate, all transported via a set of specialized transporters in the mitochondrial inner membrane.

Reducing equivalents generated by the TCA cycle, as well by glycolysis, glycogenolysis, and fatty acid oxidation drive the redox-coupled proton pumps of the respiratory chain, generating the proton motive potential that, in turn, represents the driving force for oxidative ATP synthesis. The transfer of redox potential from the cytosol to the mitochondria—via the malate aspartate shuttle and/or the glyceral-3-phosphate shuttle systems—is necessary to sustain oxidative glycolysis and/or lactate uptake and oxidation [3]. Thus, the capacity of the respiratory chain to oxidize NADH generated by glycolysis governs the myocardial physiological capacity for carbohydrate consumption.

Metabolic interactions between the matrix and cytosol are also important in pathological conditions, such as heart failure and ischemia/reperfusion injury. During ischemia, an accumulation of NADH and succinate occurs, creating an excessively reduced system that affects both matrix and cytosolic metabolic reactions. There is conflicting evidence on the route of succinate accumulation within the mitochondria during ischemia [4, 5]. However, the detrimental effects of excessive succinate accumulation impairing tissue salvage are not disputed. NAD(P)(H) redox state is hypothesized to influence the metabolic transitions during ischemia because the TCA cycle, substrate transport, and glycolytic thermodynamics and kinetics are strongly influenced by matrix NAD(P)(H) redox state [6]. Upon reperfusion, respiration on pathologically high concentrations of succinate contributes to ischemia-reperfusion injury through the production of pathological ROS levels [4]. Mitochondrial respiration on succinate is also known to result in elevated matrix concentrations of oxaloacetate (OAA) that inhibit succinate dehydrogenase (SDH) in the ETC [7, 8]. It has been hypothesized that mitochondrial inner membrane potential influences the accumulation and clearance of OAA. However, a mechanistic reasoning behind this observation is lacking [9]. Regardless, a clearance route for OAA is needed to restore physiological respiration after succinate accumulation following ischemia/reperfusion.

The overarching goal of this study is to quantitatively probe the interactions between substrate transport processes, TCA cycle kinetics, redox state, and oxidative ATP synthesis in suspensions of purified cardiac mitochondrial under experimental conditions designed to: (1.) characterize the kinetics of transport processes; (2.) elucidate a systems-level understanding of TCA cycle kinetics driven by different substrate conditions; (3.) test hypothesized pathways associated with the metabolic response to ischemia and reperfusion; and (4.) identify a theoretical/computational framework for simulating the interplay of these processes under a variety physiological and pathological conditions. The kinetic model of mitochondrial metabolism identified in this study is built from appropriately thermodynamically constrained rate equations for the substrate transporters, TCA cycle enzymes, electron transport component, and oxidative phosphorylation components, illustrated in Figure 1. Several prior models have integrated many of these components [10-18], with the model of Saito et al. [18] representing a particularly comprehensive integration of most of the components illustrated in Figure 1. However, since the Saito et al. model is identified primarily from quasi-steady data it is not intended to capture the time-course kinetics analyzed here.

**Figure 1:**
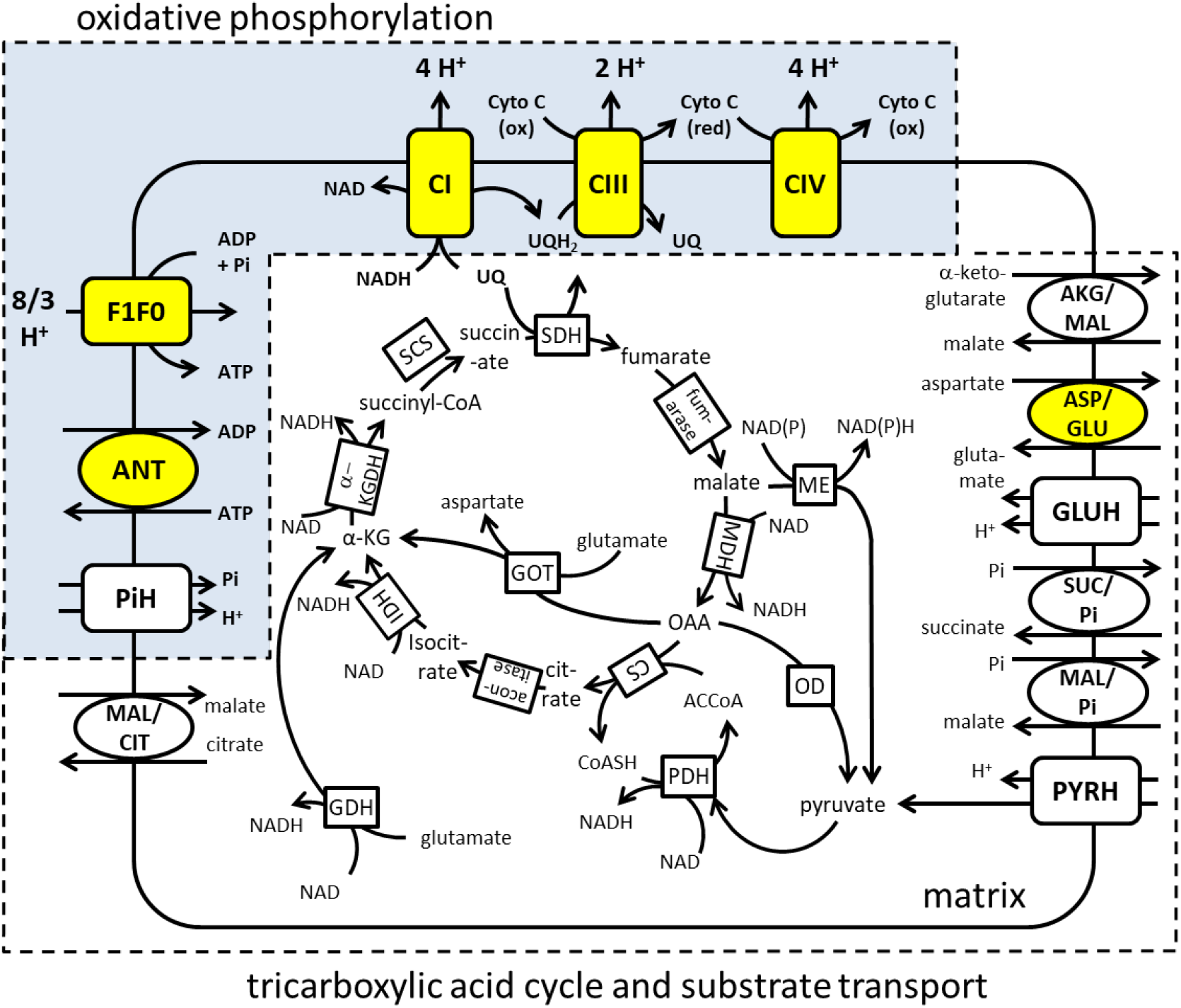
Model diagram. This diagram shows the components included in the computational model of mitochondrial metabolism. Transporters highlighted in yellow indicate electrogenic transport. Abbreviations: inorganic phosphate (Pi), glutamate oxaloacetate transaminase (GOT), pyruvate dehydrogenase (PDH), citrate synthase (CS), aconitase (ACO), isocitrate dehydrogenase (IDH), glutamate dehydrogenase (GDH), alpha ketoglutarate dehydrogenase (α-KGDH), succinyl-CoA synthetase (SCS), succinate dehydrogenase (SDH), fumarase (FH), malate dehydrogenase (MDH), malic enzyme (ME), oxaloacetate decarboxylase (OD), adenine nucleotide translocator (ANT), ATP synthase (F1F0), complex 1 (CI), complex 3 (CIII), complex 4 (CIV), acetyl coenzyme A (ACCoA), coenzyme A (CoASH), ubiquinone (UQ), cytochrome C (Cyto C), oxidized (ox), reduced (red).

In this work model parameters are identified using transient respiration and relative NAD(P)H measurements from isolated rat cardiac mitochondria given various combinations and concentrations of TCA cycle substrates. Comparisons of model simulations to experimental observations yield several new hypotheses that were subsequently tested by additional experiments. For example, this cycle of model-based experimental design and experimental interrogation revealed novel insights into ROS-mediated leak and the kinetic regulation of pyruvate dehydrogenase (PDH) activity. Specifically, interpretation of experimental data using the identified computational model indicates that:

1. PDH is deactivated via phosphorylation during leak-state (state 2) respiration. Upon the initiation of oxidative phosphorylation, PDH activation occurs causing PDH activity to increase with a time constant on the order of a minute. Fitting model simulations to experiments using pyruvate and malate as substrates facilitated the identification of a novel model of the PDH phosphorylation-dephosphorylation cycle kinetics.
2. Respiration under highly reduced conditions associated with pathologically high succinate concentrations results in an elevation of cation leak conductivity compared to respiration under physiological substrate conditions. High levels of ROS are hypothesized to activate uncoupling protein (UCP) and thereby increase leak current and leak-state respiration rate under these conditions.
3. Addition of ADP to initiate oxidative phosphorylation (state 3) with pathological succinate concentrations causes a rapid reduction in respiration attributed to the accumulation of OAA resulting in the inhibition of succinate dehydrogenase (SDH).
4. Clearance of OAA during state 3 respiration on succinate is carried out relative slowly through generation of pyruvate by malic enzyme (ME) and oxaloacetate decarboxylase (OD) activity. The clearance of OAA is enhanced by the activity of glutamate oxaloacetate transaminase (GOT) in the presence of glutamate.
5. Succinate accumulation under anoxic conditions in a suspension of purified mitochondria occurs due to the reversal of SDH.

## Methods

### Mitochondrial Isolation

All protocols involving animals conformed to the National Institutes of Health Guide for the Care and Use of Laboratory Animals and were approved by the University of Michigan Animal Research Committee. Male Sprague Dawley rats, ages 10-22 weeks, were anesthetized with 100 mg/kg ketamine and 0.5 mg/kg dexmedetomidine by an intraperitoneal (IP) injection followed by 1,000 U/kg heparin delivered by the same method. The heart was cannulated via the aorta and perfused for 5 minutes with a cold cardioplegia solution (100 mM NaCl [Sigma S-9888], 25 mM KCl [Sigma P-4504], 25 mM MOPS [Sigma M-3183], 10 mM Dextrose [Sigma D-9559], 1 mM EGTA [Sigma E-4378], pH 7.2). The ventricles were isolated on ice by removing the atria and major vessels then cut vertically to make two approximately equal pieces. Each half was processed individually on ice at the same time described as follows. The tissue was minced for 3 minutes before the addition of 11 mL of 0.273 mg/mL proteinase [Sigma P-8038] in isolation buffer (IB) (200 mM mannitol [Sigma M-9647], 60 mM sucrose [Sigma S-7903], 5 mM KH_2_PO_4_ [Sigma P-5379], 5 mM MOPS [Sigma M-3183], 1 mM EGTA [Sigma E-4378], pH 7.2). The minced tissue was homogenized for < 3 minutes using a 30 mL glass homogenizer [Potter-Elvehjem Tissue Grinder, Thermo 50-365-338] before the addition of 25 mL IB with 0.1% w/w essentially fatty acid free BSA [Sigma A-6003] containing 20 *μ*L protease inhibitor cocktail [Sigma 539134] to prevent further protease activity. The homogenate was then centrifuged at 8000 *x g* for 10 minutes followed by removal of the supernatant and resuspension in IB with 0.1% w/w BSA. This step was repeated once. Next, the homogenate was centrifuged at 700 *x g* for 5 minutes. The supernatant was collected and mixed with IB with 0.1% w/w BSA followed by an 8000 *x g* centrifugation for 10 minutes. The supernatant was discarded and the pellet was resuspended in IB with 0.1% w/w BSA. Total protein content of the isolate was measured by a Quick Start Bradford Protein Assay [Bio-Rad 5000201]. Citrate synthase (CS) activity was measured using a CS assay modified from a previously published protocol [19]. Briefly, 3 *μ*L of mitochondrial isolate was diluted to 0.1 mg protein/mL in 100 mM Tris-HCl pH 7.0 [Sigma T-1503]. A final amount of 2 ug isolate protein (20 *μ*L of 0.1 mg/mL) was added to a 1 mL cuvette containing 980 *μ*L of a solution composed of 300 mM Tris-HCl pH 8.1 [Sigma T-1503], 0.25% Triton X-100 [Sigma T-8532], 0.31 mM acetyl CoA [Sigma ACOA-RO 10101893001], 0.1 mM DTNB [Sigma D-218200], and 0.5 mM oxaloacetate [Sigma O-4126]. Changes in absorbance at 412 nm over time were measured immediately after addition.

### Respirometry

Transient and steady-state mitochondrial respiration data were collected at 37°C in respiration buffer (RB) (90 mM KCl [Sigma P-4504], 50 mM MOPS [Sigma M-1254], 1 mM EGTA [Sigma E-4378], and 0.1% w/w BSA [Sigma A-6003], pH 7.2). Mitochondria were suspended in RB in a high resolution closed two-chamber respirometer (Oxygraph 2K Respirometer, Oroboros Instruments GmbH, Innsbruck, Austria) where oxygen concentration was measured every 2 seconds and oxygen flux was calculated by the DatLab software. The functional integrity of each mitochondrial isolate preparation was determined by the respiratory control ratio (RCR) defined as the ratio of steady-state oxygen flux during leak state and oxidative phosphorylation respiration (15.7 ± 0.2 [Ave±SEM]). Leak state and oxidative phosphorylation respiration were assessed under the following constituent concentrations: 5 mM KH_2_PO_4_ [Sigma P-5379], 1.5 mM MgCl_2_ [Sigma M-3634], 5 mM NaCl [Sigma S-9888], 5 mM sodium pyruvate [Sigma P-8574], 1 mM potassium L-malate pH 7.0 [Sigma M-1000], 1 mM potassium ADP [Sigma A-5285]. The percentage of mitochondria having a compromised outer membrane was calculated by the percent increase in steady-state respiration from state 3 to after the addition of 10 *μ*M cytochrome C [Sigma C-2037] (5.3% ± 0.2% [Ave ± SEM]). Based on the CS assay, mitochondrial isolate was added to each chamber to achieve a concentration of 1.17 ± 0.04 U CS/mL (Ave ± SEM) in the Oxygraph for each experimental run. For each experimental run, depending on the substrates provided to the mitochondria, NaCl was added to achieve a final sodium concentration of 10 mM. Experiments were started with the addition of NaCl, if needed, and KH_2_PO_4_ followed by mitochondria. After 130 seconds, substrate(s) were added to the chamber initiating state-2 leak. Potassium ADP was added 150 seconds after substrates. Oxygen flux is reported as an average of biological replicates ± the standard error of the mean except for the 5 mM succinate ± 1 mM or 5 mM glutamate experiments. These are shown as one biological replicate because of the variability of the rise in respiration towards the end of the experimental run. Anoxia-reoxygenation experiments were carried out by adding 10 mM NaCl, 1 mM pyruvate, 0.5 mM malate, 2 mM ADP, and 2.5 mM KH_2_PO_4_ to the chamber followed by mitochondria. A timer was started once the oxygen concentration in the chamber reached zero to indicate the start of anoxia. The stopper of the chamber was removed to reoxygenate the system. To collect samples, the stir bar was stopped and 1 mL or 1.5 mL of suspension was extracted from the chamber with a syringe. For the collection during anoxia, at 0, 2, 4, 6, 10, and 15 minutes, mitochondria were extracted with a syringe flushed with deoxygenated RB (sparged with N_2_). Reoxygenation samples, collected at 16, 17, 20, 25 minutes, were collected with a syringe flushed with RB.

### NAD(P)H Autofluorescence

Mitochondria were suspended in 1 mL RB in 24-well black assay plates [4titude 4ti-0262] at the same concentration as in the respirometry experiments determined by the CS assay. NAD(P)H autofluorescence was measured from the top of each well at 37°C in a modular multi-mode microplate reader with an automated injection system [BioTek Synergy H1, Agilent]. The excitation and emission wavelengths were set to 340 nm and 460 nm, respectively. The gain and read height were automatically adjusted by the Gen5 3.11 software to optimize fluorescence measurements of mitochondria suspended in RB in the presence of 2.5 mM KH_2_PO_4_, 5 mM NaCl, 0.5 mM sodium pyruvate, 0.25 mM potassium malate pH 7.0, and 10 *μ*M rotenone [Sigma R-8875]. The gain was set to 134 and the read height 6.75 mm. Only 4 wells were used at once to increase the number of data point collected during each run. For all runs, fluorescence in the 4 wells was measured every 6 seconds with double orbital mixing in between measurements. Experiments were run in duplicate and the raw fluorescence data were averaged before analysis. For each experimental run, depending on the substrates provided to the mitochondria, NaCl was added to achieve a final sodium concentration of 10 mM. Mitochondria were added to 1 mL of RB containing NaCl, if needed, and KH_2_PO_4_, mixed manually, bubbles were removed, and fluorescence was measured over 2 minutes and 10 seconds. Substrate(s) were automatically injected and the plate was mixed for 5 seconds before measuring redox during leak state for 2 minutes and 30 seconds. After the automatic addition of potassium ADP and 5 seconds of mixing, oxidative phosphorylation state was measured for 5 minutes. To measure the fully reduced state, 10 *μ*M of rotenone was added manually, the plate was mixed for 5 seconds, and fluorescence was measured for 3 minutes. In runs where pyruvate and malate were not already present, 0.5 mM sodium pyruvate and 0.25 mM potassium malate pH 7.0 were added with rotenone to ensure a fully reduced state. Relative NAD(P)H was calculated using state 1 (only mitochondria and phosphate present) as the fully oxidized state and the presence of rotenone as the fully reduced state, therefore:

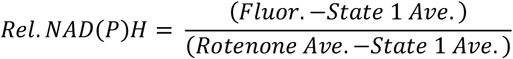

Relative NAD(P)H is reported as an average of biological replicates ± the standard error of the mean unless stated otherwise.

### Metabolite Extraction

Suspensions of mitochondria (1 mL or 1.5 mL) were collected from the respirometer chamber after stopping the stir bar and immediately placed in 200 uL of 3 N perchloric acid [244252, Sigma] with 3 mM EDTA [E4884, Sigma]. All solutions added to the sample were kept ice cold. The sample was vortexed and spun at 15,000g for 5 minutes at 4°C. The supernatant was then transferred to 60 uL of 10 N KOH, vortexed, and neutralized with 2 N KOH with 40 mM TES [T6541, Sigma] and 300 mM KCl [P4504, Sigma] and/or with 2 N perchloric acid with 2 mM EDTA. The precipitate was pelleted at 15,000 x g for 10 minutes at 4°C and the supernatant was aliquoted for several assays and stored at −80°C.

### Pyruvate, Malate, Alpha Ketoglutarate, ATP, ADP, AMP, and Succinate Measurements

Pyruvate, ADP, AMP, and succinate measurements were based on the following enzymatic reactions:

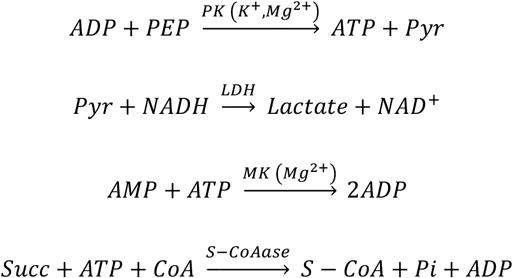

These reactions were measured via NADH absorbance at 340 nm at room temperature. Extracted sample was mixed with a solution containing 278 mM TEA [90279, Sigma], 38.2 mM MgSO_4_ [M7506; Sigma], 152.8 mM KCl [P4504, Sigma], 1.39 mM phosphoenolpyruvate [PEP; AAB2035806; VWR], and 120 µg of NADH [Calzyme Laboratories, San Luis Obispo, CA]. The pyruvate reaction was initiated with the addition of 8.25 U lactate dehydrogenase [LDH; L2500; Sigma].Once pyruvate was depleted, the ADP reaction was catalyzed by adding 4.7 U pyruvate kinase [PK; P1506; Sigma]. Once all ADP is consumed, the AMP reaction is catalyzed by adding 7.2 U of myokinase [MK; M3003; Sigma] and 0.075 mM ATP [A2383, Sigma]. After AMP was depleted, the succinate reaction was catalyzed by adding 506 mU of succinyl-CoA synthetase [S-CoAase; E-SCOAS, Megazyme, Lansing, MI], 0.3 mM Coenzyme A [CoA; 234101, Calbiochem], and 0.075 mM ATP [A2383, Sigma].

The assay for ATP measurement was previously published and followed [20].

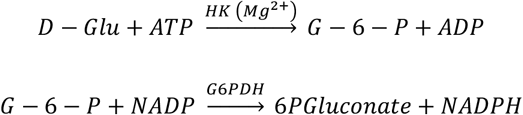

This reaction was measured via NADPH absorbance at 340 nm at room temperature. Extracted sample was mixed with an assay mixture (2X-1 ug/uL NADP+ [Calzyme Laboratories, San Luis Obispo, CA], 56.8 mM Tris-HCl [T1503, Sigma] with 5 mM MgCl_2_ solution [M0250, Sigma], and 1 U/uL Glucose-6-Phosphate Dehydrogenase [G8404, Sigma]). The reaction was catalyzed with the addition of 10 mM glucose [D9559, Sigma] and 1 U hexokinase [HK; H4502, Sigma].

### Computational Model

The model used to simulate and analyze experiments on suspension of purified mitochondria integrates the processes diagrammed in Figure 1 The model takes the form of 69 ordinary differential equations describing kinetics of biochemical reactants, mitochondrial inner membrane potential, cation (Mg^2+^, K^+^) concentrations, and pH. The model formulation is based on an existing simulation framework [15, 21-23] and integrates numerous enzyme and transporter component models from prior studies [18, 24-35]. The model is described in detail in the Appendix.

Adjustable parameters in the model are largely associated with enzyme, transporter, pump, and exchanger activities/maximal fluxes. In addition, several new model components, including models of the pyruvate dehydrogenase phosphorylation-dephosphorylation cycle, the dicarboxylate carrier, and the oxoglutarate carrier were developed and parameterized by matching model simulations to observed data.

### Data Availability Statement

Computer codes and associated data are available at: https://github.com/beards-lab/mitochondrial-carb-tca-oxphos.

## Results

Mitochondria were isolated from rat cardiac ventricular tissue and oxygen consumption rate and NAD(P)H autofluorescence were measured over time. Mitochondria were added to the respirometer chamber at time zero. Substrates were added causing mitochondria enter a leak state (state 2) where NAD(P)H is produced via the TCA cycle and a relatively low level of respiration is sustained, called the leak state. Oxidative phosphorylation (state 3) is initiated upon the addition of ADP. Under our experimental conditions, most of the ADP is phosphorylated to ATP, and the mitochondria enter another leak state termed state 4 following oxidative phosphorylation. The respiratory rate of this state is affected by the level of contaminating ATPases present in various degrees of all isolated mitochondrial preparations. This experimental procedure was repeated with a variety of substrate combinations detailed below. Model simulations were fitted to the data from condition informed identification of parameters in the model illustrated in Figure 1.

### Mitochondrial Metabolism with Pyruvate and Malate Substrates

Figure 2 illustrates results from experiments with pyruvate (0.5 mM initial concentration) and malate (0.25 mM initial concentration) as substrates. Figure 2A and C show results obtained using an initial buffer inorganic phosphate (Pi) concentration of 0.5 mM. Figure 2B and D show results using initial phosphate levels of 2.5 mM. During the initial leak state (state 2), a low level of oxygen consumption is sustained with approximately 80% of the total NAD(P)(H) pool reduced under both concentrations of Pi. After ADP addition (*t* = 280 seconds), oxidative phosphorylation (state 3) is initiated resulting in an increase in respiration and a drop in NADH as it is oxidized by complex I of the ETC. Under limiting phosphate conditions, oxygen consumption rate reaches a maximum of 50 nmol O_2_·min^-1^·UCS^-1^ and gradually decays over time. As the oxygen consumption rate decays during oxidative phosphorylation, the relative level of NAD(P)H increases. When ADP is limiting, as under 2.5 mM initial Pi, respiration rate increases to a maximal level of approximately 80 nmol O_2_·min^-1^·UCS^-1^ after at approximately 45 seconds after addition of ADP. The 0.375 mM added ADP is consumed after approximately 120 seconds of state-3 respiration and the system returns to a final leak state (state 4) with high NAD(P)H. By adjusting parameters associated with the subset of enzymes highlighted in Figure 2E, the model captures both oxygen flux and changes in redox state under both phosphate concentrations. The model fit to these data is not highly sensitive to the activities of enzymes downstream of isocitrate dehydrogenase (IDH) in the TCA cycle because, as illustrated below, under these experimental conditions the alpha-ketoglutarate (α-KG) formed by IDH is largely transported out of the matrix into the buffer.

**Figure 2:**
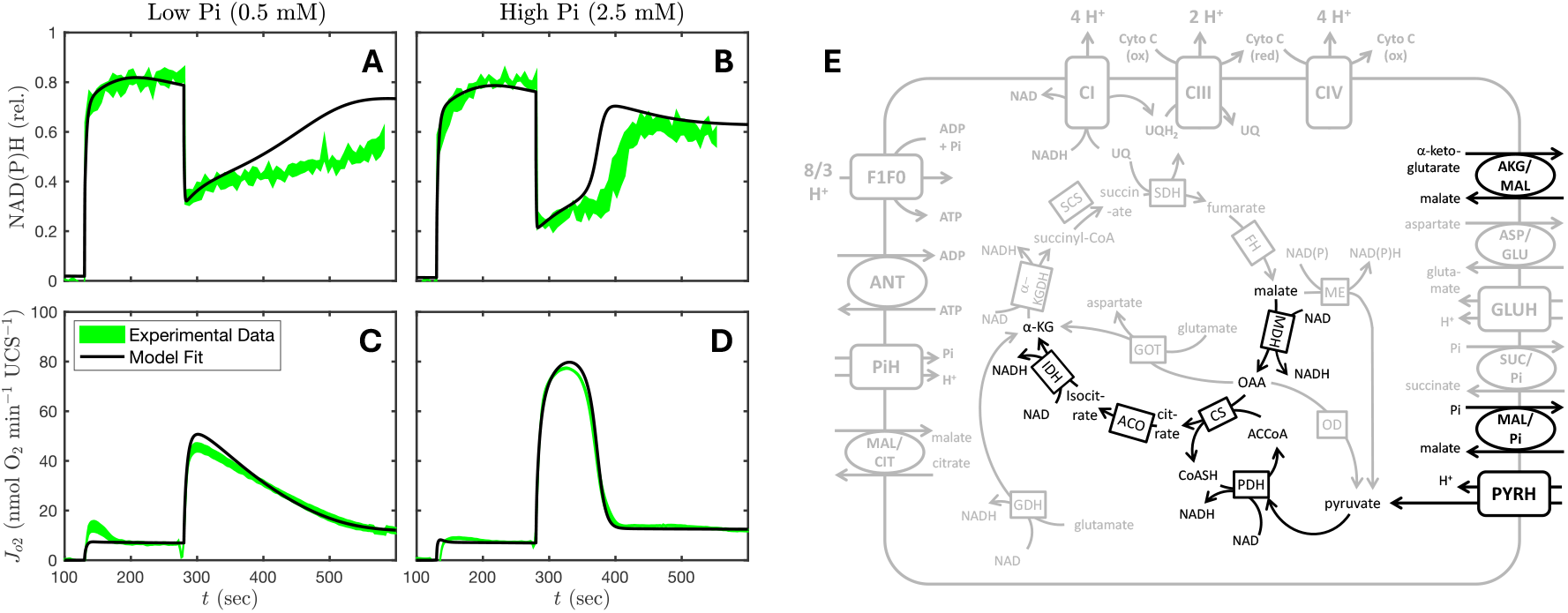
Respiration on 0.5 mM pyruvate and 0.25 mM malate with 0.375 mM ADP. A) and B) show the relative NAD(P)H time courses at 0.5 mM and 2.5 mM inorganic phosphate (Pi). C) and D) show oxygen consumption rate at 0.5 mM and 2.5 mM Pi. Substrates are added at time t = 130 seconds, initiating leak state respiration. Oxidative phosphorylation is initiated by addition of ADP at t = 280 seconds. The model (black curves) accurately captures the kinetics of oxygen consumption and redox state during pyruvate and malate respiration in vitro. E) The subset of model components to which model predictions are most sensitive for the simulation of this experiment are highlighted.

Data on metabolite concentrations of the pyruvate and malate experiments are compared to model simulations in Figure 3. The figure shows measurements of ATP, ADP, AMP (A and B), pyruvate, malate, and α-KG (C and D), and α-KG and succinate (E and F) under the experimental conditions described for the results in Figure 2. Since metabolic measurements are obtained from biochemical quenching and extracting the experimental preparation at discrete time points, data are compared to model simulations of total concentration in the suspension. These total concentrations are nearly identical to buffer concentrations since mitochondrial are diluted in an appropriately 1:1000 ratio of mitochondrial-to-buffer volume. Data and model simulations indicate under conditions of low and high Pi, approximately three quarters (75%) of oxidized pyruvate generates α-KG that exits the mitochondrial matrix rather than be oxidized by alpha-ketoglutarate dehydrogenase (α-KGDH). This finding indicates that α-KGDH contributes little to NAD(P)H production during pyruvate and malate-fueled respiration under these conditions. In support of this, less 10 µM of succinate was measured experimentally at all time points. As expected for adenine nucleotides, ADP is phosphorylated to ATP during the oxidative phosphorylation state, with a faster and more complete phosphorylation occurring under high phosphate conditions compared to limiting phosphate conditions. Experiments also reveal low levels of AMP (ranging from 2 to 6 µM following oxidative phosphorylation), indicating adenylate kinase activity that may be associated with contamination in the mitochondrial isolate. As with the oxygen flux and redox state, the model matches the metabolite flux under both phosphate concentrations, illustrating the model’s ability to capture the dynamics responsible for NAD(P)H and ATP production.

**Figure 3:**
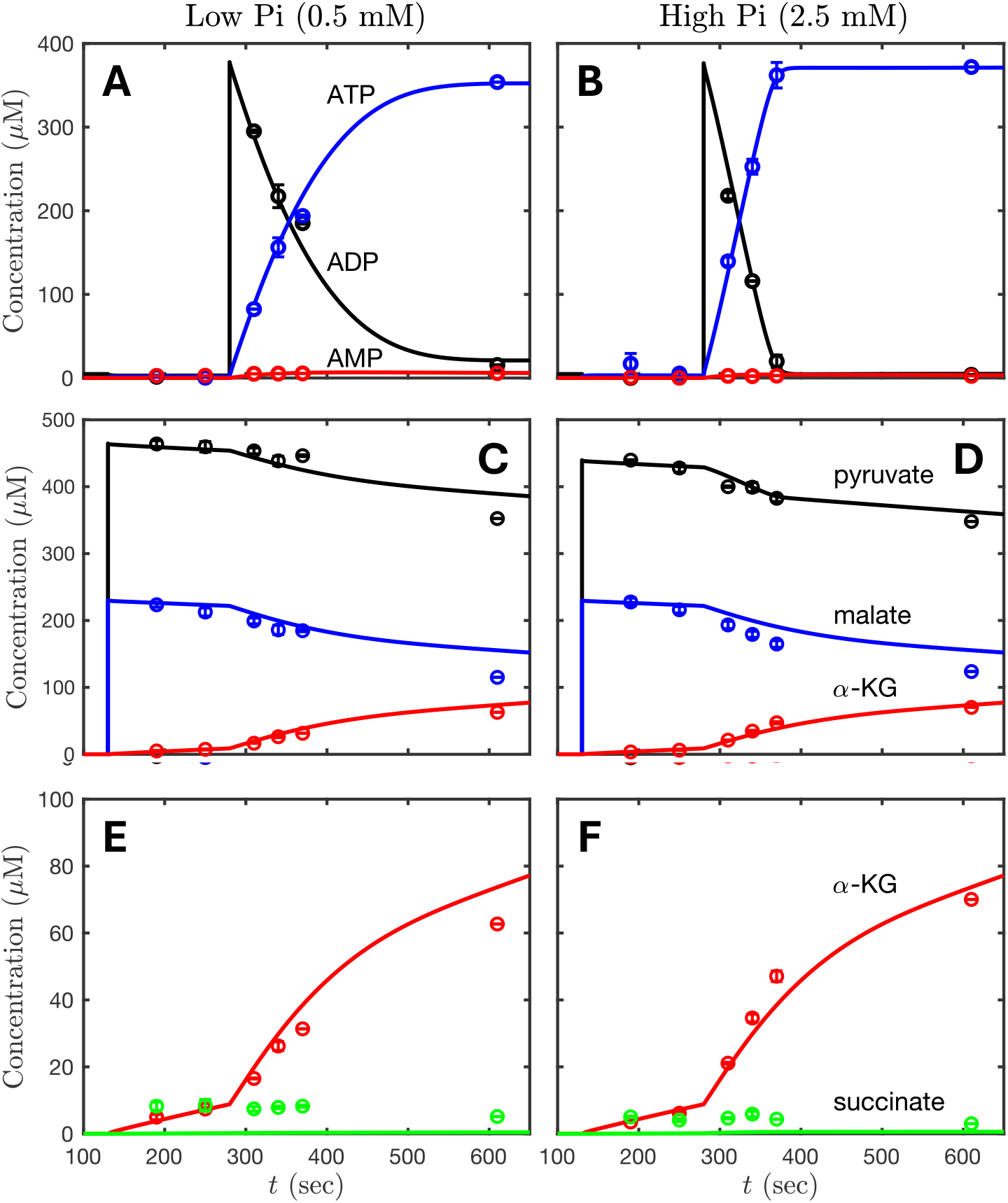
Kinetics of ATP, ADP, AMP (A and B), pyruvate, malate, and alpha ketoglutarate (α-KG) (C and D), and α-KG and succinate (E and F) during respiration on 0.5 mM pyruvate and 0.25 mM malate with 0.5 mM Pi and 2.5 mM Pi. Data and associated model predictions represent total concentrations of reactants in buffer containing suspensions of mitochondria for the experimental protocol described in Figure 2. The majority of pyruvate consumed is exported as α-KG out of the matrix into the buffer under these experimental conditions.

### Kinetics of PDH Phosphorylation and Dephosphorylation

Analogous experiments with pyruvate and malate substrates were performed at a non-limiting ADP concentration of 1 mM. Results are shown in Figure 4. The addition of 1 mM ADP results in a higher rate of respiration compared to the maximal rate obtained with 0.375 mM ADP. Under this high ADP condition, for the high Pi case, the respirometer chamber runs out of oxygen around *t* = 500 seconds, before all of the ADP is phosphorylated (Figure 4B). While the rate of oxygen consumption reaches a peak value approximately 45 seconds after the additions of 0.375 mM ADP, the peak of respiration rate after 1 mM ADP addition is not reached until nearly 60 seconds later.

**Figure 4:**
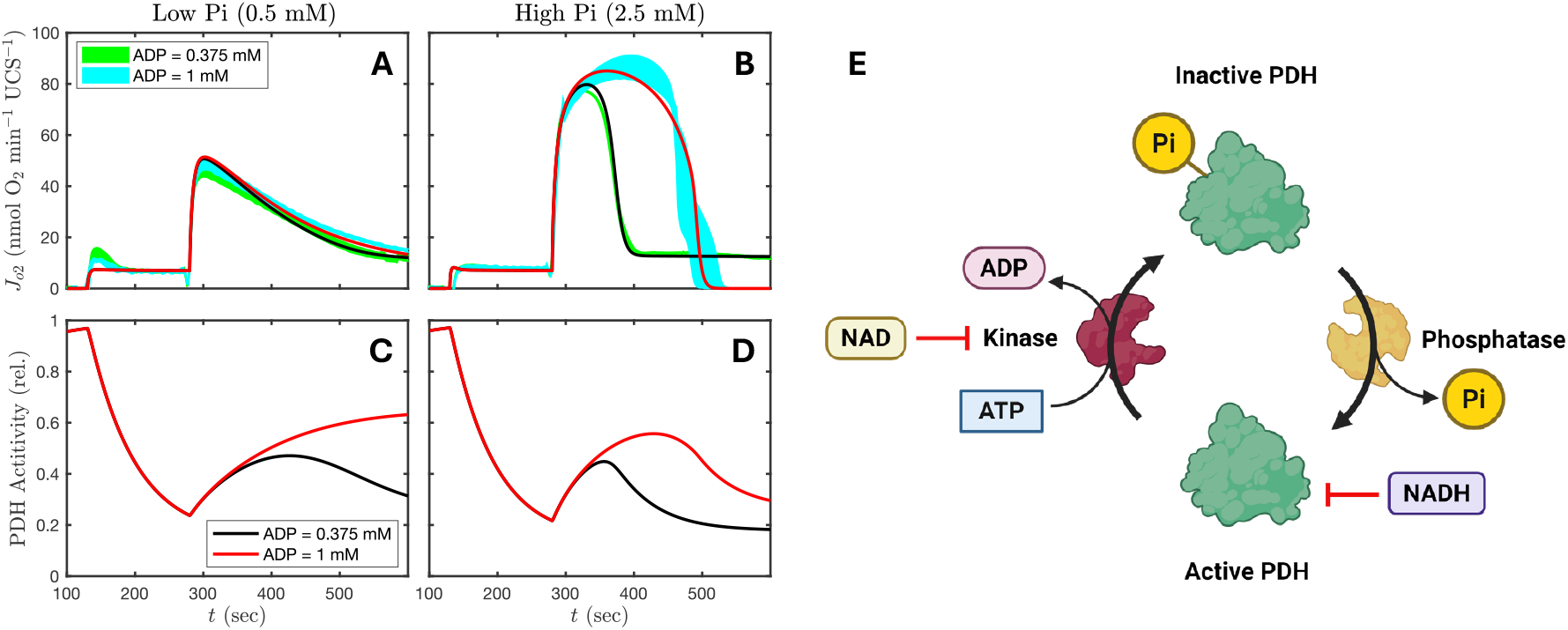
Kinetics of pyruvate dehydrogenase (PDH) deactivation and activation. Panels A and B show oxygen consumption time courses during respiration on 0.5 mM pyruvate and 0.25 mM malate with 0.375 mM and 1 mM ADP with 0.5 mM Pi and 2.5 mM Pi. Panels C and D show model-predicted PDH activity, with phosphorylation (deactivation) occurring during the initial leak state, and dephosphorylation (deactivation) following initiation of oxidative phosphorylation. E) PDH regulation is governed by NAD(H) redox level and ATP availability. Figure made with Biorender.

The model is able to match the observed slow increase in respiratory rate during oxidative phosphorylation for both the limiting and non-limiting ADP cases based on a model of PDH activation/deactivation governed by matrix ATP and NAD(H). This regulatory system responds to the energetic and metabolic status of the matrix allowing mitochondria to alter PDH activity in response to the needs of the cell in vivo [36, 37]. The model simulates the relative proportion of two states for the PDH enzymes: a phosphorylated (inactive) and dephosphorylated (active) state. The phosphorylation process requires ATP and is inhibited by NAD. The dephosphorylation process is inhibited by NADH (Figure 4E). The resulting predicted PDH activity time courses associated with the model fits to the data are shown in Figure 4C and D. Model simulations assume that the initial condition for PDH is the active dephosphorylated state, consistent with a lack of ATP and redox potential. Under reduced conditions associated with state-2 leak, PDH is gradually inactivated by phosphorylation. After the additions of ADP, when NAD(P)H drops, predicted PDH activity begins to increase due to dephosphorylation. The level of activation acts as a rate limiting step for oxygen consumption on pyruvate and malate substrates. With a higher concentration of ADP, a higher proportion of PDH is predicted to become active, allowing for a higher rate of respiration. The time it takes to activate PDH determines the time at which maximum oxygen flux is reached.

The hypothesized role PDH phosphorylation/dephosphorylation cycle governing state-3 respiratory kinetics with pyruvate substrate emerges from fitting the model to the data shown in Figures 2 and 4. Two ways to independently test this hypothesis are to vary the time between substrate and ADP addition and to reverse the order of addition of ADP and Pi substrates in the experiment. Figure 5A shows experimental data obtained from varying the length of the leak state (state 2). As the duration of the leak state decreases, there is less time for PDH to become deactivated. Therefore, the model predicts a higher maximal rate of respiration during oxidative phosphorylation (state 3) and an earlier peak in respiration compared to longer-duration leak-state simulation. The model predictions match the trends seen in the experimental data (Figure 5B). Figure 5C shows experimental results of reversing the order of addition of ADP and Pi substrates. Here the initial experimental buffer contains ADP but not Pi, and oxidative phosphorylation is initiated by the addition of Pi. When ADP is supplied in the absence of Pi, there is no matrix ATP available to phosphorylate PDH. Thus, the model predicts that PDH remains essentially 100% active. Upon addition of Pi, to initiate oxidative phosphorylation, a maximal peak in respiration is reached much more quickly than in the control conditions. The model predictions match these data (Figure 5D). These results support the hypothesis that PDH phosphorylation/dephosphorylation kinetics governs respiration with pyruvate and malate and validate the kinetic model of PDH phosphorylation/dephosphorylation.

**Figure 5:**
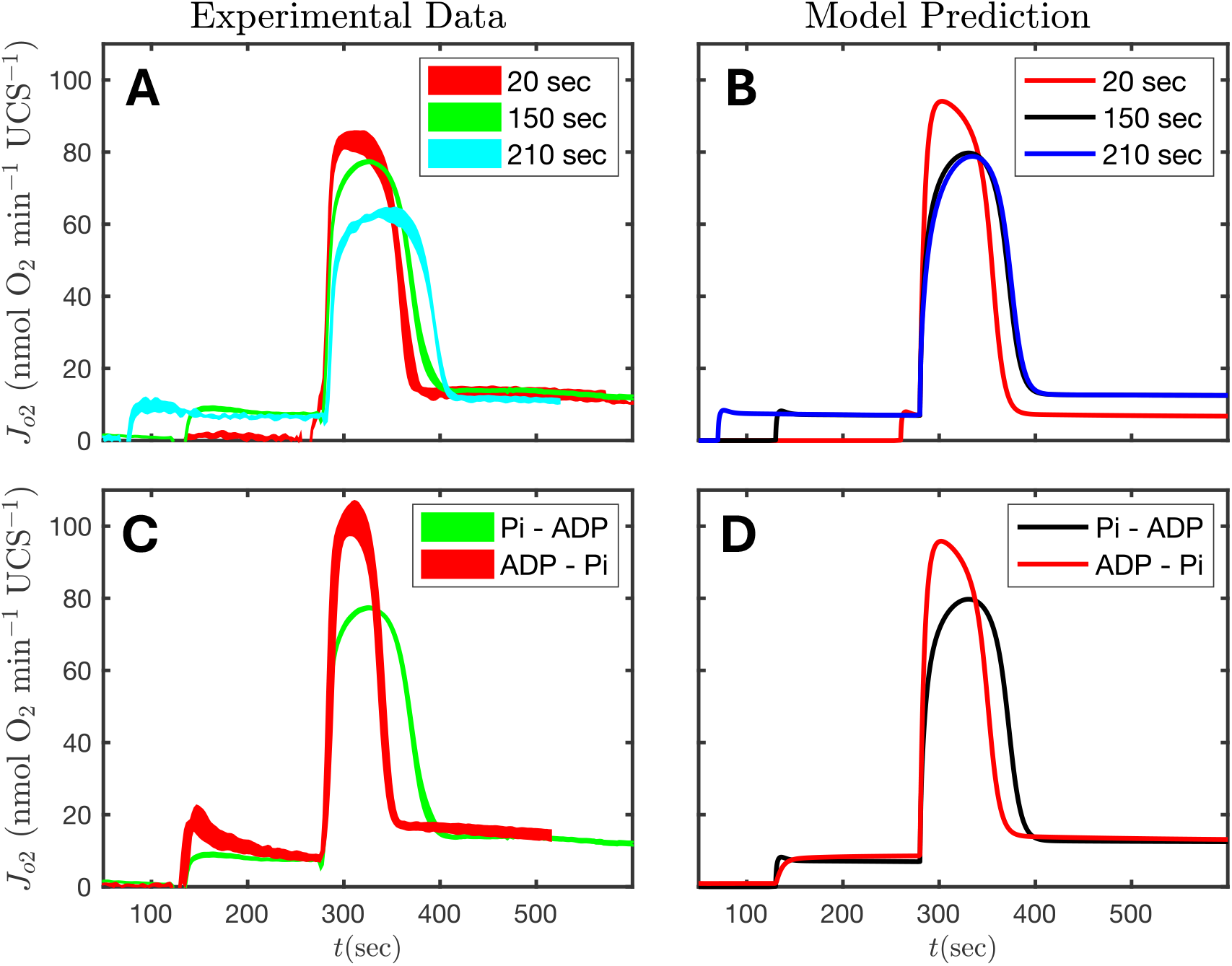
Validation of the pyruvate dehydrogenase (PDH) regulation model. Experiment with pyruvate and malate substrates were conducted to slow the rate of inhibition of PDH during the initial leak state. A) The duration of the leak state was varied, resulting in a faster increase in respiration following addition of ADP at t = 280 seconds. B) Model-predictions of the experiments illustrated in panel A match the experimental observations. C) In one set of experiments, labeled ‘ADP - Pi’, ADP was supplied without phosphate during the initial leak state and oxidative phosphorylation was initiated by the addition of phosphate at t = 280 seconds. The ‘ADP - Pi’ protocol is compared to the standard ‘Pi – ADP’ protocol in which ADP is added to initiate oxidative phosphorylation. D) Model-predictions of the experiments illustrated in panel C match the experimental observations.

### Mitochondrial Metabolism with Limiting Pyruvate

To investigate oxidative ATP synthesis under limiting pyruvate, the protocol of Figure 2 was repeated with 0.05 mM initial pyruvate and 0.25 mM initial malate. Results from this protocol are shown in Figure 6. As diagrammed in Figure 2, we expect 3 NADH molecules to be generated for each pyruvate consumed under these experimental conditions. Moreover each NADH drives the pumping of 10 charges across the inner membrane. Since approximately one ATP is synthesized for each 11/3 charges translocated across the membrane, nearly all of the initial pyruvate is expected to be consumed in phosphorylation 0.375 mM of ADP. Thus, maximal respiration is lower in these conditions compared to the high-pyruvate conditions of Figure 2. NADH recovers at the end of the oxidative phosphorylation state, reaching a peak at approximately *t* = 460 seconds. Subsequently NADH drops as the available pyruvate is exhausted.

**Figure 6:**
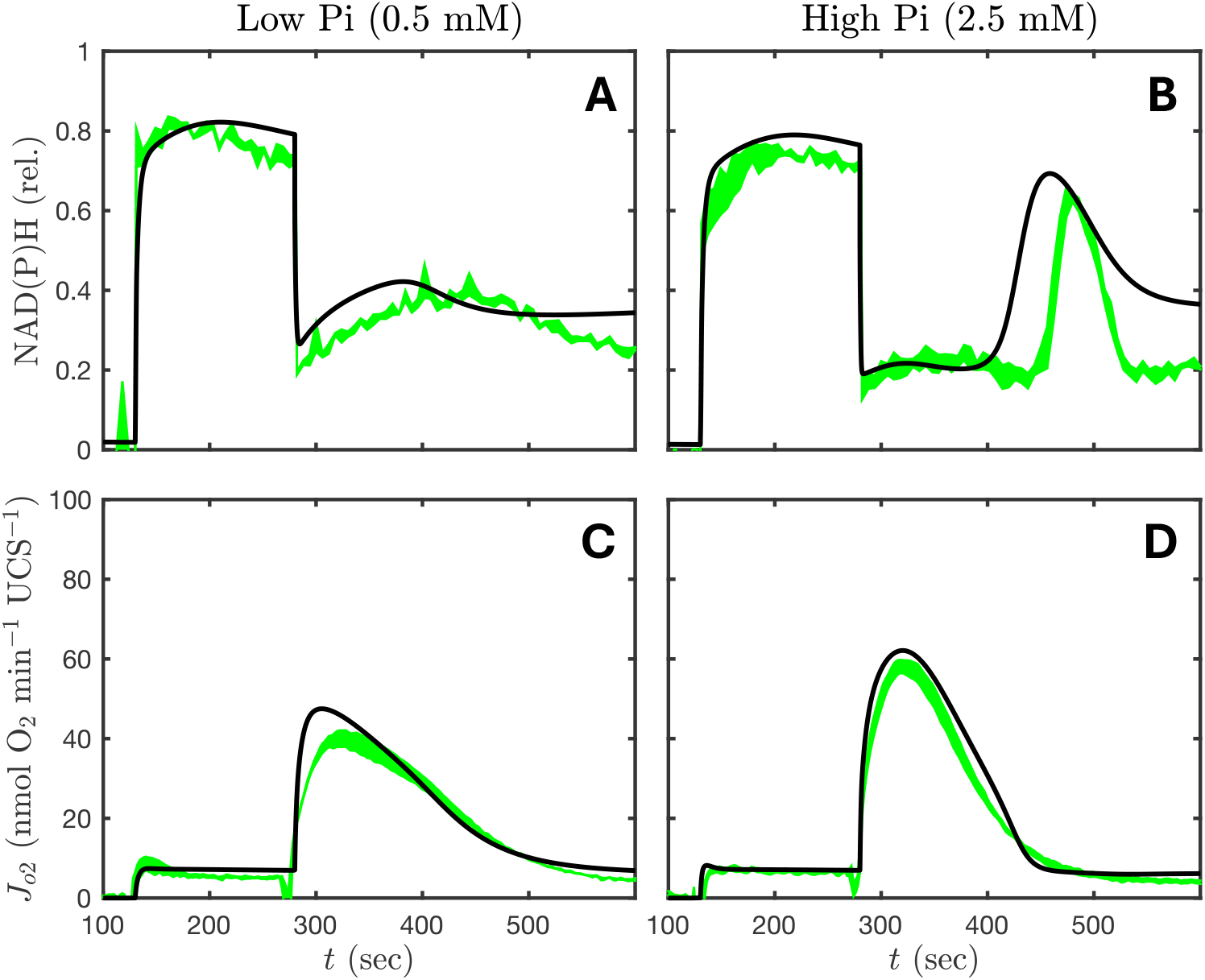
Respiration on limiting (0.05 mM) pyruvate and 0.25 mM malate with 0.375 mM ADP. A) and B) show the relative NAD(P)H time courses at 0.5 mM and 2.5 mM inorganic phosphate (Pi). C) and D) show oxygen consumption rate at 0.5 mM and 2.5 mM Pi. Substrates are added at time t = 130 seconds, initiating leak state respiration. Oxidative phosphorylation is initiated by addition of ADP at t = 280 seconds. The model (black curves) accurately captures the kinetics of oxygen consumption and redox state during pyruvate and malate respiration in vitro.

### Mitochondrial Metabolism with α-Ketoglutarate and Malate Substrates

Data and model simulations for experiments measuring oxygen consumption and relative redox state was measured with 5 mM α-KG and 0.25 mM malate added to initiate the leak state (state 2) are shown in Figure 7. In Figures 7A and B, the initial leak state NAD(P)H reaches a level approximately half of what is observed with pyruvate and malate substrates. The respiration curves (Figures 7C and D) show behavior similar to pyruvate and malate-fueled respiration with a 25% lower maximal oxygen consumption rate. Model simulations of this experiment are sensitive to the activities of α-KGDH and downstream enzymes highlighted in Figure 7E. The subset of enzymes and transporters interrogated during α-KG and malate-fueled respiration makes up the other half of the TCA cycle compared to respiration on pyruvate and malate (Figure 7E).

**Figure 7:**
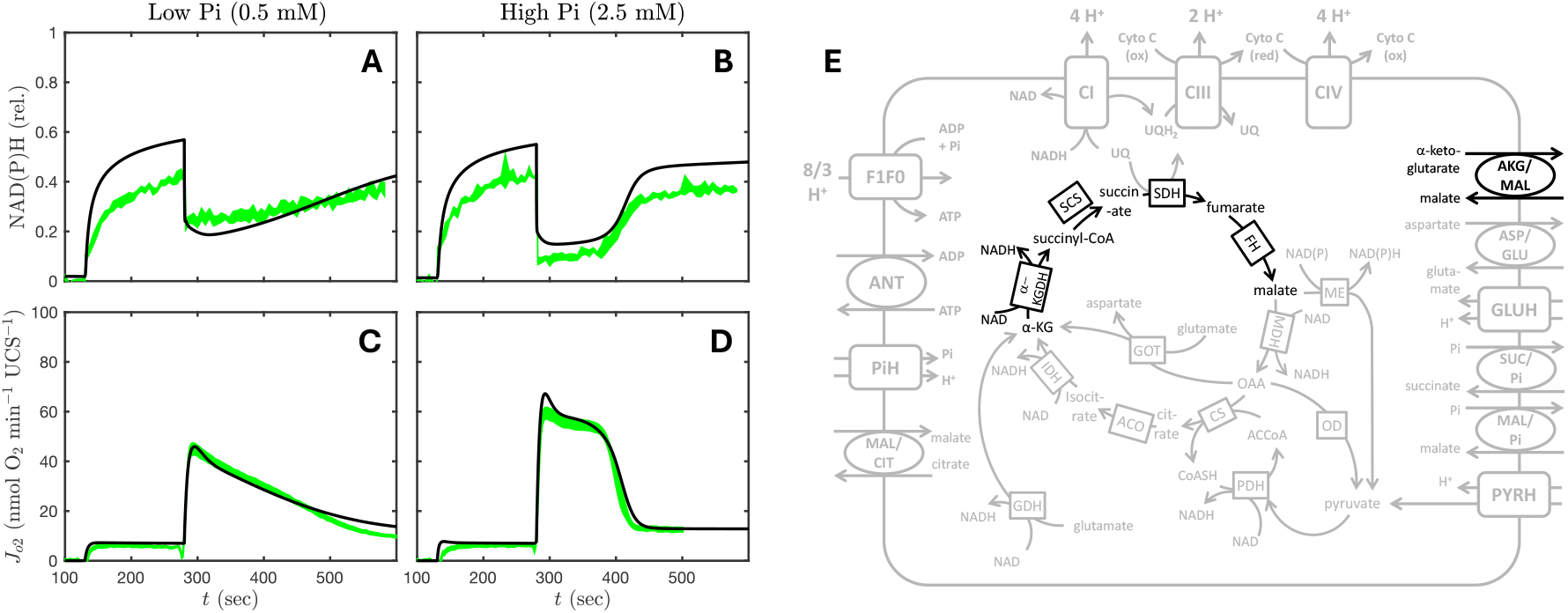
Respiration on 5.0 mM α-ketoglutarate (α-KG) and 0.25 mM malate. A) and B) show the relative NAD(P)H time courses at 0.5 mM and 2.5 mM inorganic phosphate (Pi). C) and D) show oxygen consumption rate at 0.5 mM and 2.5 mM Pi. Substrates are added at time t = 130 seconds, initiating leak state respiration. Oxidative phosphorylation is initiated by addition of ADP at t = 280 seconds. The model (black curves) accurately captures the kinetics of oxygen consumption and redox state during α-KG and malate respiration in vitro. E) The subset of model components to which model predictions are most sensitive for the simulation of this experiment are highlighted.

### Mitochondrial Metabolism with Succinate Substrate

Figure 8 shows relative redox state (A and B) and oxygen consumption (C and D) kinetics obtained for respiration on pathologically high concentrations of succinate (5 mM initial concentration). Here the observed behavior is quantitatively different from that observed under pyruvate and malate as well as α-KG and malate substrate conditions. The observed leak-state respiration is approximately 3-fold greater than that observed under pyruvate and malate conditions and the NAD(P)H reaches a level of effectively 100% reduced. Upon ADP addition, there is a rapid drop in NAD(P)(H) and a transient spike in respiration. The state-3 spike in respiration is not associated with a sustained oxidative phosphorylation state. Rather, oxygen consumption slowly rises following the initial spike, reaching a peak around *t* = 500 seconds for the low phosphate case and around *t* = 600 seconds for the high phosphate case. Note that since the precise timing of late peaks in oxygen consumption varies between experimental replicates, averaging across replicates introduces smoothing artefacts that obscure the true nature of this feature. Thus, the data shown in Figure 7C and D are from a single representative experiment.

**Figure 8:**
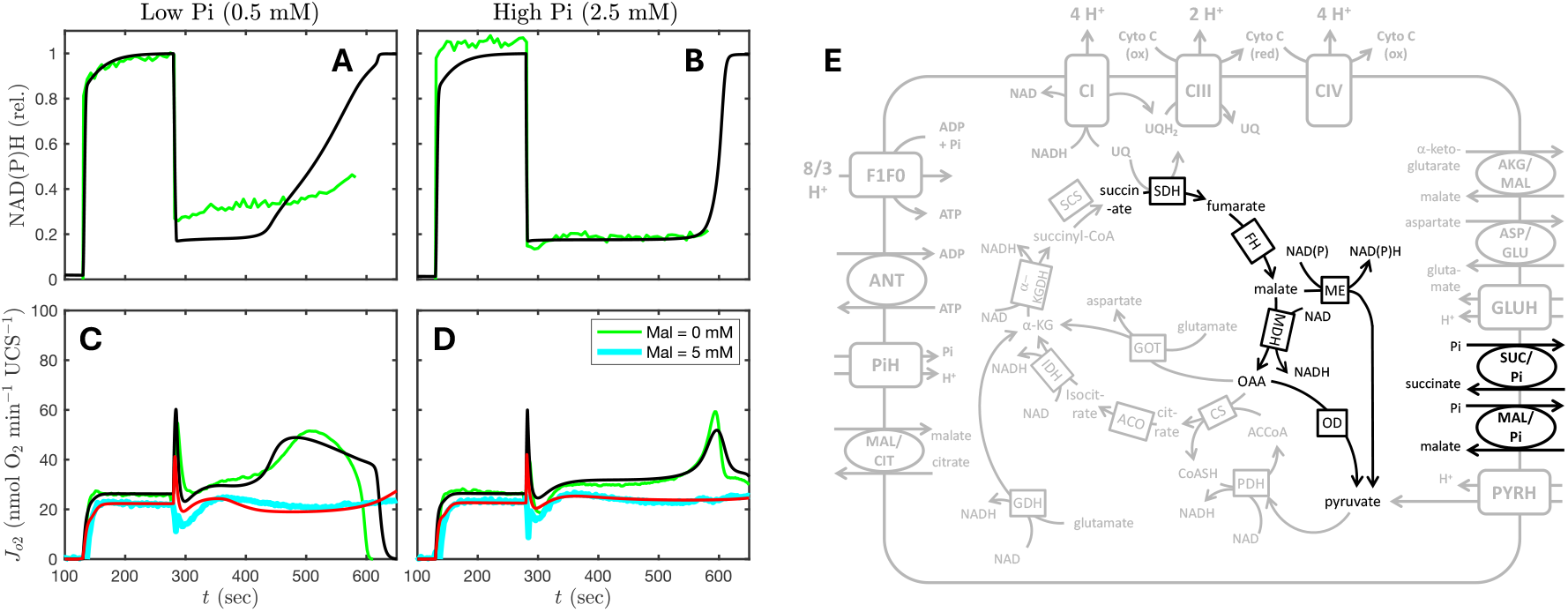
Respiration on 5.0 mM succinate with and without 5.0 mM malate. A) and B) show the relative NAD(P)H time courses at 0.5 mM and 2.5 mM inorganic phosphate (Pi). C) and D) show oxygen consumption rate at 0.5 mM and 2.5 mM Pi. The high oxygen leak during leak-state respiration is predicted to be caused by relatively high membrane potential, the lower proton-pumping efficiency of succinate-fueled respiration compared to complex-I substrates, and a ROS-activated increase in leak. An inhibition of oxidative phosphorylation is predicted to be caused by the rapid production of oxaloacetate (OAA), which inhibits succinate dehydrogenase, by malate dehydrogenase upon the addition of ADP. The gradual rise in respiration following the addition of ADP at t = 280 seconds is predicted to be driven by generation of pyruvate by malic enzyme and oxaloacetate decarboxylase activity. The addition of 5 mM malate (yellow and red) further inhibits oxidative phosphorylation due increased OAA production. E) The subset of model components to which model predictions are most sensitive for the simulation of this experiment are highlighted.

Model-based analysis of these data predicts the observed high leak rate is due to (1) elevated membrane potential compared to that obtained with respiration on complex I substrate, (2) a lower number of hydrogen ions pumped across the inner membrane per one oxygen atom reduced compared to complex I respiration, and (3) increased leak conductivity. The increased leak conductivity is hypothesized to arise from activation of ROS-sensitive uncoupling protein in the inner mitochondrial membrane [38, 39]. Furthermore, the model predicts that the reduction in respiration following the initial spike associated with addition of ADP is due to inhibition of succinate dehydrogenase (SDH) by oxaloacetate (OAA). Model-predicted time courses of malate and OAA for these experiments are shown in Figure 9A and B for each Pi concentration. The relatively high NAD(P)H level obtained during leak is predicted to inhibit malate dehydrogenase (MDH), resulting in a build-up of malate concentrations between 5-7 mM in the matrix (Figure 9). The rapid oxidation of NADH after ADP is added to the system allows MDH to generate high concentrations of OAA from this accumulated malate. The OAA, predicted to transiently reach matrix concentrations of approximately 0.2 mM, inhibits SDH causing a depression in oxidative phosphorylation respiration. Eventually, oxygen consumption rises under both low- and high-Pi conditions, predicted to follow from the clearance of OAA by citrate synthase and oxaloacetate decarboxylase (OD). Malic enzyme (ME) is predicted to provide pyruvate to fuel the generation of acetyl-CoA to help clear the accumulated OAA. Experiments using 5 mM initial malate in combination with 5 mM succinate to test the mechanisms of OAA-mediated inhibition of SDH and OD- and ME-mediated clearance of OAA are discussed below.

**Figure 9:**
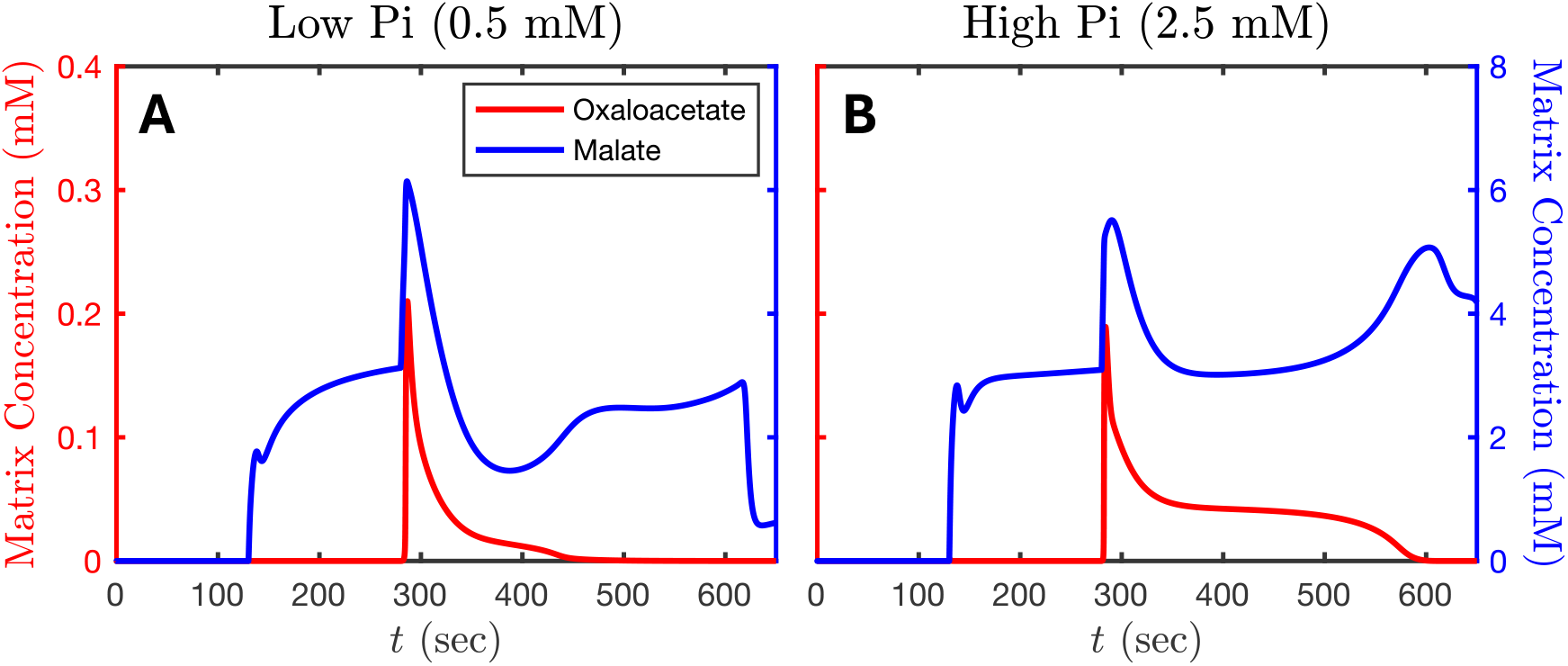
Model prediction of matrix malate and oxaloacetate fluxes during succinate-driven respiration at low (A) and high (B) inorganic phosphate (Pi) concentrations. Malate and oxaloacetate concentrations rise upon the addition of ADP at t = 280 seconds and is predicted to be responsible for the observed inhibition of oxidative phosphorylation. The clearance of oxaloacetate corresponds to the rise in oxygen consumption rate during succinate respiration (Figure 8).

### Mitochondrial Metabolism with Malate Substrate

The predicted activities of ME and OD associated with the succinate-fueled respiration experiment led to the hypothesis that mitochondria could respire on malate alone, with ME generating pyruvate to fuel complex I respiration. Results of applying the same protocol used in Figure 2 with 5 mM initial malate as the sole substrate are shown in Figure 10. Under these conditions mitochondria achieve a oxidative phosphorylation respiration rate of approximately 20 O_2_·min^-1^·UCS^-1^ under both Pi concentrations, approximately one quarter of the rate achieved with high Pi with pyruvate and malate substrates shown in Figure 2. These data support the hypothesis that the activity of ME can support malate-only driven respiration.

**Figure 10:**
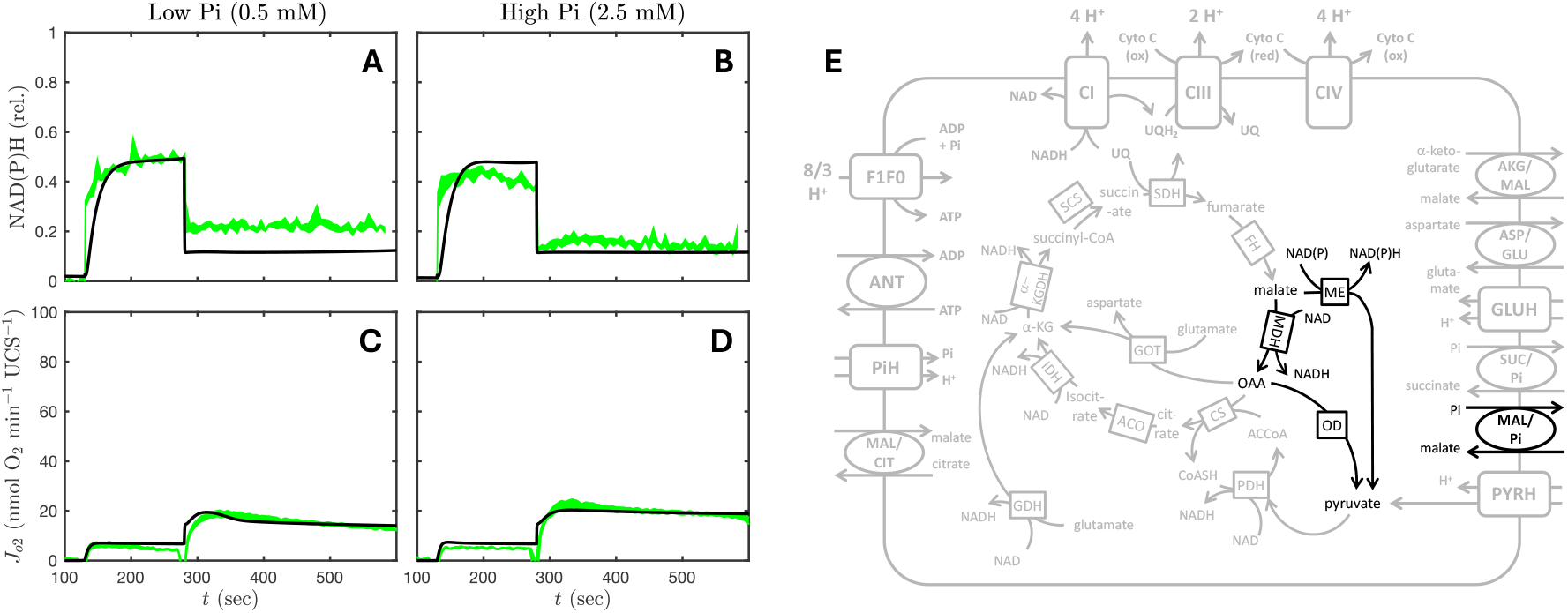
Respiration on 5.0 mM malate with 0.375 mM ADP. A) and B) show the relative NAD(P)H time courses at 0.5 mM and 2.5 mM inorganic phosphate (Pi). C) and D) show oxygen consumption rate at 0.5 mM and 2.5 mM Pi. Substrates are added at time t = 130 seconds, initiating leak state respiration. Oxidative phosphorylation is initiated by addition of ADP at t = 280 seconds. Model simulations are shown as black curves. E) The subset of model components to which model predictions are most sensitive for the simulation of this experiment are highlighted.

### Testing Model Predictions Under Oxygenated Conditions

The role of malate and OAA build-up in inhibiting SDH during succinate-driven metabolism was tested by the addition of 5 mM malate under succinate substrate condition. Experimental data (blue curves) and the model predictions (red curves) are shown in Figure 8C and D. Note that these model predictions do not represent a fit to the data but rather a prediction of the identified model. The model predicts that the addition of 5 mM exogenous malate represses oxygen consumption rate via increased OAA concentration compared to respiration on succinate alone. The fidelity of model predictions compared to data supports the interpretation of the mechanism driving the kinetics of respiration on succinate.

Simulations also predict that glutamate-oxaloacetate transaminase (GOT) represents a potential route of OAA clearance under conditions of succinate-fueled respiration. Results from experiments using 5 mM succinate and glutamate (1 or 5 mM) are shown in Figure 11A and B. The inhibition of SDH by OAA is effectively abolished in the presence of 5 mM glutamate under both low- and high-Pi conditions.

**Figure 11:**
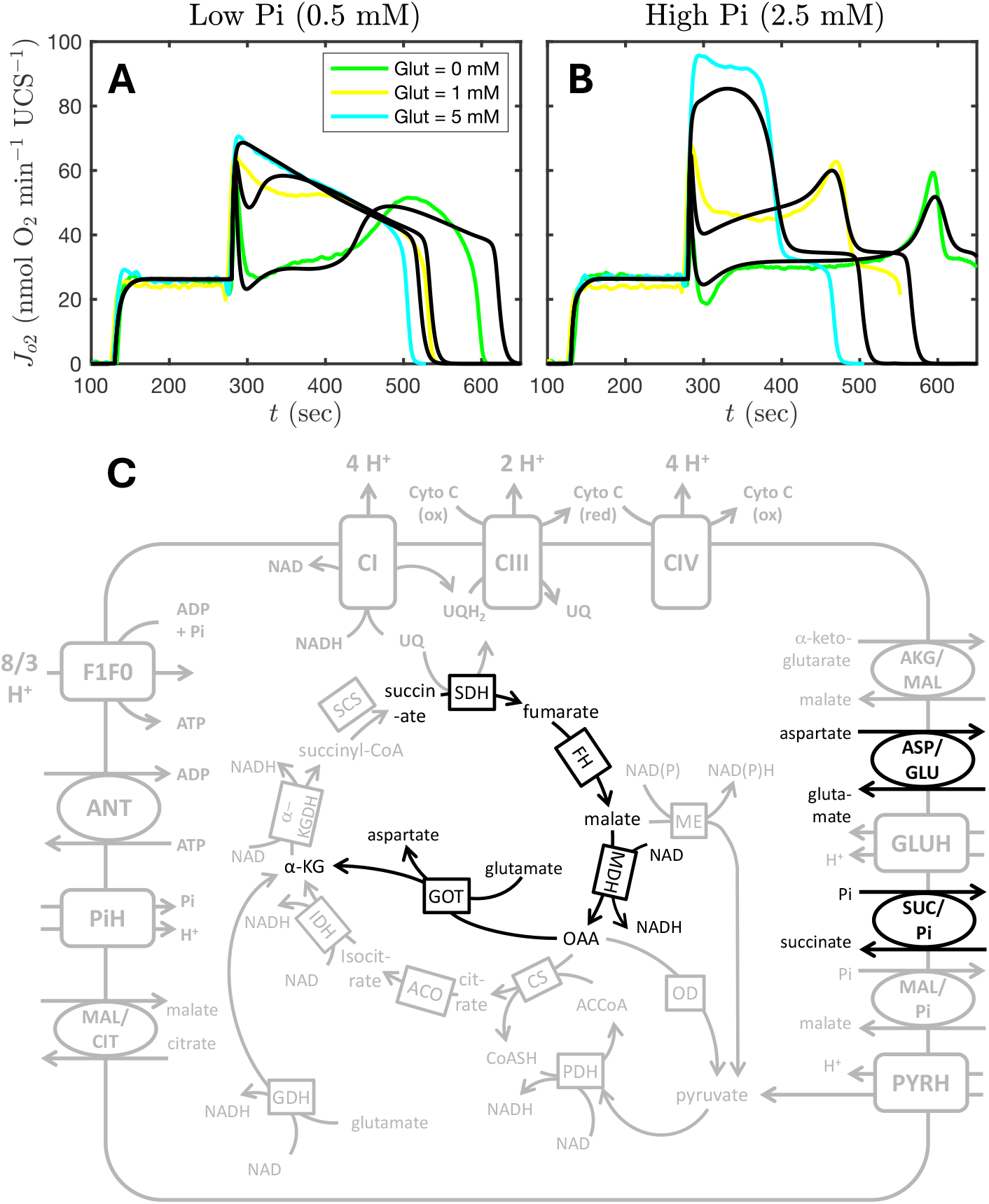
Respiration on 5.0 mM succinate ± 1 mM (purple) or 5 mM glutamate (blue) with low (A) and high (B) Pi. A) and B) show oxygen consumption rate at 0.5 mM and 2.5 mM Pi. Substrates are added at time t = 130 seconds, initiating leak state respiration. Oxidative phosphorylation is initiated by addition of ADP at t = 280 seconds. Model simulations shown as black curves. E) The subset of model components to which model predictions are most sensitive for the simulation of this experiment are highlighted.

### Quasi-steady behavior

Figure 12 shows comparisons of model simulations to data from Bazil et al. [33] and Vinnakota et al. [40]. In these experiments an ATP hydrolyzing enzyme was titrated into the buffer to achieve quasi-steady state respiration over a range of demand in vitro. NAD(P)H redox level, membrane potential (ΔΨ), cytochrome redox level, and ADP were measured with different concentration of inorganic phosphate. The model is able to effectively capture the relationships between these state variables associated with oxidative phosphorylation and rate of respiration. For experimental details see [33, 40].

**Figure 12:**
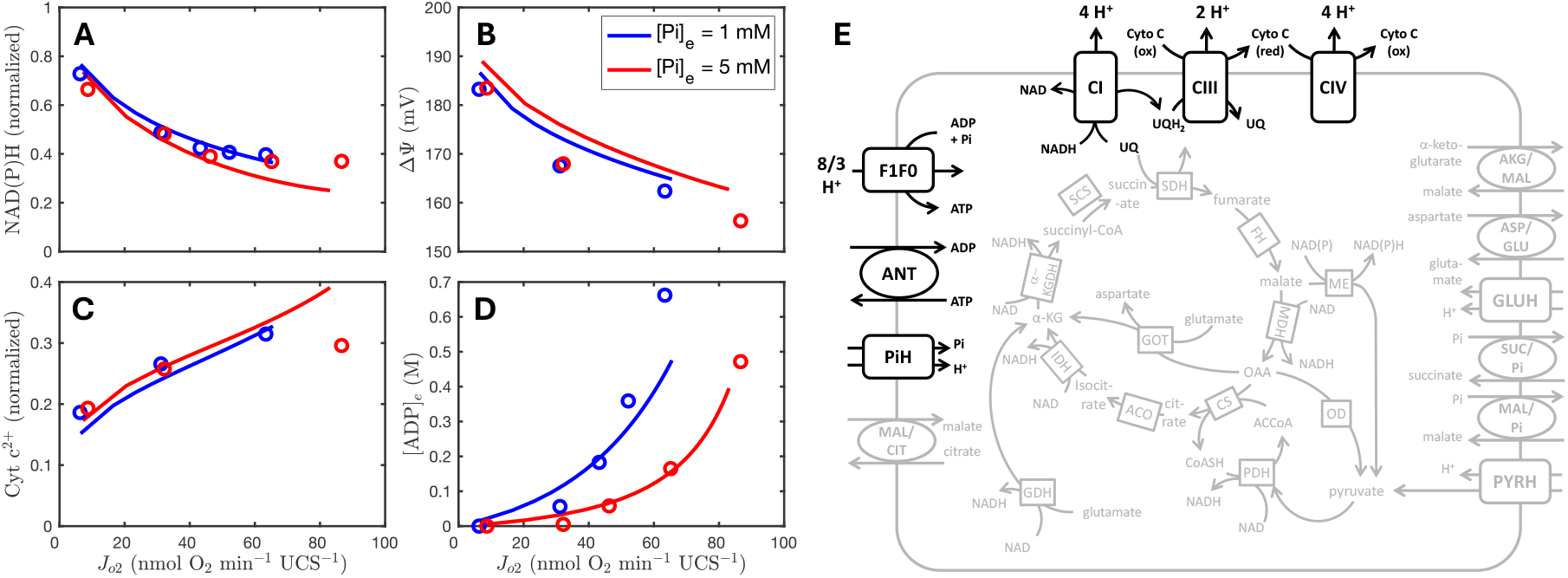
Oxidative phosphorylation under quasi-steady conditions. Experimental data are from Vinnakota et al. [40] in which quasi-steady state conditions were maintained by addition of ATP-hydrolyzing apyrase enzyme. Model simulations are compared to experimental data on NAD(P)H, membrane potential, cytochrome c redox, and ADP concentration at two concentrations of buffer inorganic phosphate. Panel E highlights the subset of model components to which model predictions are most sensitive for the simulation of this experiment.

### Simulating Anoxia and Reoxygenation

The computational model is shown to realistically represent the observed behavior of mitochondria respiring on a variety of substrates under oxygenated conditions, including under conditions of pathologically elevated succinate.

To probe mitochondrial metabolism in ischemia and reperfusion conditions, our experimental system and associated computational model were exposed to anoxia and reoxygenation. Fluxes of oxygen consumption, pyruvate, malate, ATP, ADP, AMP, and succinate for the anoxia-reoxygenation experiments are shown in Figure 13. For this experiment, 2 mM ADP was added to the chamber to ensure a long enough oxidative phosphorylation state (state 3) for oxygen to become completely depleted (Figure 13A). The time point *t* = 0 is defined here as the time at which oxygen is depleted. After 15 minutes of anoxia, the chamber was opened allowing for reoxygenation. The rate of pyruvate and malate consumption is diminished during anoxia compared to reoxygenation (Figure 13B). As expected, phosphorylation of ADP is halted at the start of anoxia resulting in a gradual drop in ATP during the anoxic phase (Figure 13C). The rise in AMP concentration accounts for the loss of ATP and the minimal flux in ADP. This result is matched well by the model prediction with the activity of adenylate kinase.

**Figure 13:**
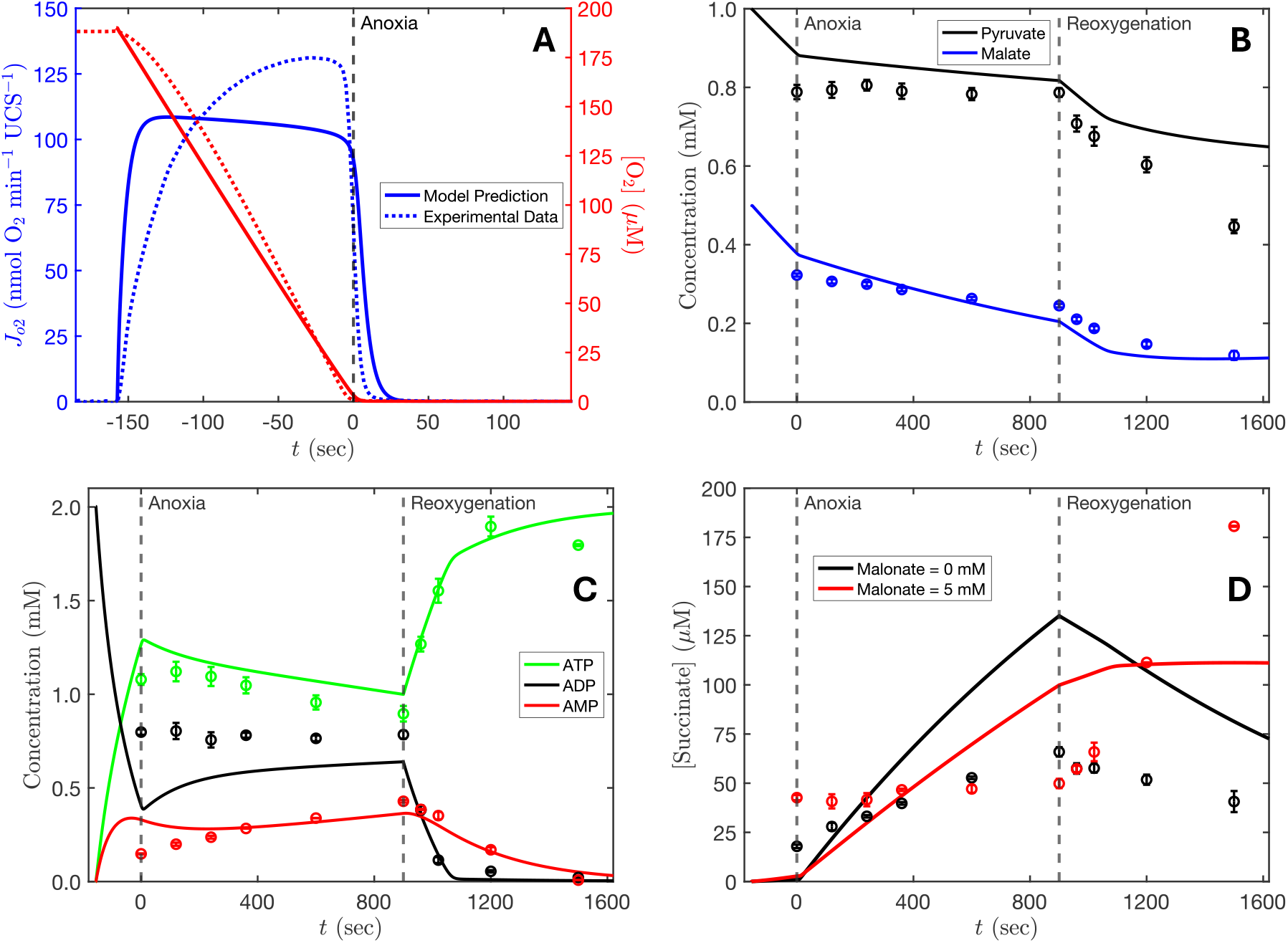
Mitochondrial metabolism in anoxia. Respiration is initiated by addition of substates (1.0 mM pyruvate, 0.5 mM malate) and 2.0 mM ADP at t = 0. Oxygen in the chamber is exhausted after approximately 155 seconds. A) Measured oxygen consumption rate (blue) and concentration (red) are compared to model simulations. B) Model simulations are compared to experimental data on pyruvate (green) and malate (purple). C) Model simulations are compared to measurements of ATP (green), ADP (black), and AMP (red) during anoxia and reoxygenation. ATP decreases over the period of anoxia while AMP increases and ADP stay relatively constant. Upon reoxygenation, ADP is quickly phosphorylated to produce ATP and AMP is gradually depleted. D) Model simulations are compared to data on succinate during anoxia and reoxygenation. Succinate accumulation is seen during the anoxic period (black) in the data and predicted by the model. Upon reoxygenation, succinate is oxidized by SDH. In the presence of malonate (red), an inhibitor of SDH, no changes in succinate concentration are seen during the anoxic period. Succinate is elevated at time zero and continues to rise during reoxygenation with malonate due to canonical TCA activity.

Data show that at the start of anoxia, about 18 µM of succinate is present and steadily accumulates in the system to a concentration of approximately 70 µM after 15 minutes (Figure 13D). During the reoxygenation period, succinate is consumed at roughly the same rate it is produced during anoxia. Model simulations are qualitatively consistent with observed data, although the predicted rate of succinate accumulation is roughly twice as fast than observed experimentally.

Malonate, an inhibitor of SDH, was used to test the primary route of succinate synthesis. With malonate present, succinate rises to a higher level during the initial oxygenated phase, presumably because it competes with succinate binding to SDH and slows its oxidation rate. The addition of malonate also prevents the rise in succinate during the anoxia period. Finally, with malonate added, succinate continues to build up during the reoxygenation period. These data suggest that the primary route of succinate accumulation is reversal of the SDH reaction during the anoxia period.

## Discussion

We have conducted a panel of experimental assays and associated computer simulations to probe the kinetics of mitochondrial ATP synthesis in suspensions of purified cardiac mitochondria. By identifying and validating a model of mitochondrial metabolism by fitting simulations to data from each experiment, hypotheses were developed that explained the behavior seen in oxygen consumption and relative NAD(P)H transients. Taken together, respiration and NAD(P)(H) data from the experiments with pyruvate and malate (Figures 2-4, 6), α-KG and malate (Figure 7), succinate (Figure 8), and malate (Figure 10) provide a rich data set to identify the activities of the enzymes and transporters in the model illustrated in Figure 1. The identified model was then used to design experiments to test several key model predictions, as illustrated in Figures 5, 8, and 13.

Results from pyruvate and malate-driven metabolism experiments (Figures 4 and 5) highlighted novel insights to the regulation of PDH activity, which plays an important role in governing cellular energetics [41] and represents a potentially important etiological driver in diabetes and certain cancers [42-45]. Validation experiments, varying the length of the leak state on the kinetics of oxidative phosphorylation state oxygen consumption rate (Figure 5A and B), are effectively predicted by the model. In addition, the model accurately predicts the effects of interchanging the additions ADP and inorganic phosphate (Figure 5C and D). When mitochondria are incubated with ADP during the leak state instead of inorganic phosphate, PDH cannot be phosphorylated. Thus, the observed rapid rise in oxygen consumption upon the initiation of oxidative phosphorylation is predicted to be achieved with maximally activated PDH.

The finding that the majority of α-KG synthesized by mitochondria respiring on pyruvate and malate is exported from the matrix, shown in Figure 3, may be explained in part by the regulation of α-KGDH by calcium and the sensitivity of α-KGDH to NADH inhibition [28, 46-48]. Calcium ion is virtually absent under the conditions of the experiments presented in this study. Under such low calcium concentrations, α-KGDH is inhibited at relatively low concentrations of NADH. Therefore, during pyruvate and malate driven respiration, the proportion of α-KG exported from the matrix is higher than what is used by α-KGDH. Although a low concentration is succinate is measured (Figure 3E and F), it remains virtually constant throughout each state.

Fitting mitochondrial respiration on pathological concentrations of succinate (Figure 8) led to the development of a number of hypotheses: In order to match the relatively high oxygen flux seen during leak state, the ion leak conductivity must be greater than it is with other substrates. The model assumes that ROS generated during respiration on succinate activates the UCP-3 cardiac uncoupling protein ROS [39, 49, 50]. Details are provided in the Appendix. Additionally, during respiration on succinate, upon the addition of ADP, inhibition of oxidative phosphorylation occurs within 10 seconds. The model predicts a rapid buildup of OAA following addition of ADP results in SDH inhibition (Figure 7). Respiration increases only when the excess OAA is cleared. The activities of ME and OD are predicted to contribute to the clearance of OAA but have slow rates when compared to GOT when glutamate is present (Figure 11).

The predicted activity of ME and OD in the succinate-fueled respiration experiments led to the hypothesis that mitochondria could sustain respiration on malate alone (Figure 9). Although oxygen flux is relatively low during oxidative phosphorylation with malate as fuel, the observations highlight the ability of ME and OD to sustain oxidative phosphorylation.

It is hypothesized that succinate accumulation during anoxia can occur via canonical TCA activity [5]. However, SDH reversal under anoxic conditions predicted by the model aligns with other studies that have also explored this question [4, 51]. The model predicts a rate of succinate accumulation in anoxia that is approximately double the observed rate (Figure 13D). This discrepancy may be due to an inability of the model—which is identified based on experiments with SDH, fumarase, and malate dehydrogenase operating in the forward reaction direction—to reproduce the reverse kinetics of these reactions.

## Conclusion

A computational model of mitochondrial ATP synthesis and substrate transport was developed and validated using oxygen consumption and relative redox data from suspension of purified mitochondria. This process revealed mechanisms underlying mitochondrial metabolism under a diverse combination of substrates and conditions that are difficult to measure experimentally, especially transiently. During pyruvate and malate driven metabolism, α-KGDH was found to contribute very little to NAD(P)H production. In addition, PDH regulation was found to determine the time it takes to reach maximum oxygen flux during oxidative phosphorylation respiration. The relatively high leak respiration seen during succinate oxidation was hypothesized to be due to high membrane potential, the ratio of ions pumped around the inner membrane for each succinate consumed, and the opening of ROS-activated UCP-3 triggered by pathological ROS production. Upon ADP addition, rapid production of toxic levels of OAA inhibits SDH and lowers oxygen flux. ME and OD activities contribute to OAA clearance and also enable a relatively low rate of respiration under conditions where malate is the only substrate. GOT was found to be an effective route of OAA clearance in the presence of glutamate. Finally, succinate accumulation during anoxia occurs via partial reversal of the TCA cycle by SDH reversal.

The developed model’s predictive capabilities highlight the potential to be used as a tool in understanding mitochondrial behavior under not only physiological but pathological conditions. This foundation has the potential to be built upon by adding models such as beta-oxidation and cytosolic reactions (ex. glycolysis [3]) that would expand the model’s predictive capabilities to metabolism at a cellular level.

## Additional Information

### Author Contributions

Contextualization: NLC, SD, JB, DAB

Methodology and formal analysis: NLC, SD, FVB, DAB

Investigation, software, visualization and data curation: NLC, DAB

Writing -- original draft: NLC, SD, DAB

Writing -- review and editing: NLC, FVB, JN, DB

Funding acquisition and supervision: DAB Resources: DAB

Project administration: DAB

All authors have approved the final version of the manuscript.

## Appendix

### A. Notation and Abbreviations

The reactants and reference species for all metabolites in the system are listed in Table A1 below. Brackets are used to indicate concentration. For example [ATP]_x_ denotes the overall concentration of all species of the reactant ATP in the matrix. For example

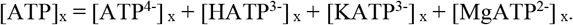

Binding polynomials are represented using *P*, where the subscript indicates the reactant. For example,

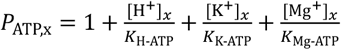

is the binding polynomial for matrix ATP and *K*_H-ATP_, *K*_K-ATP_, and *K*_Mg-ATP_ are dissociation constants.

### B. List of Model Components

This appendix lists the basic components of the model. Tables A1 and A2 list the state variables and reaction and transport fluxes included in the model. Table A3 lists fixed model parameters.

**Table 1:**
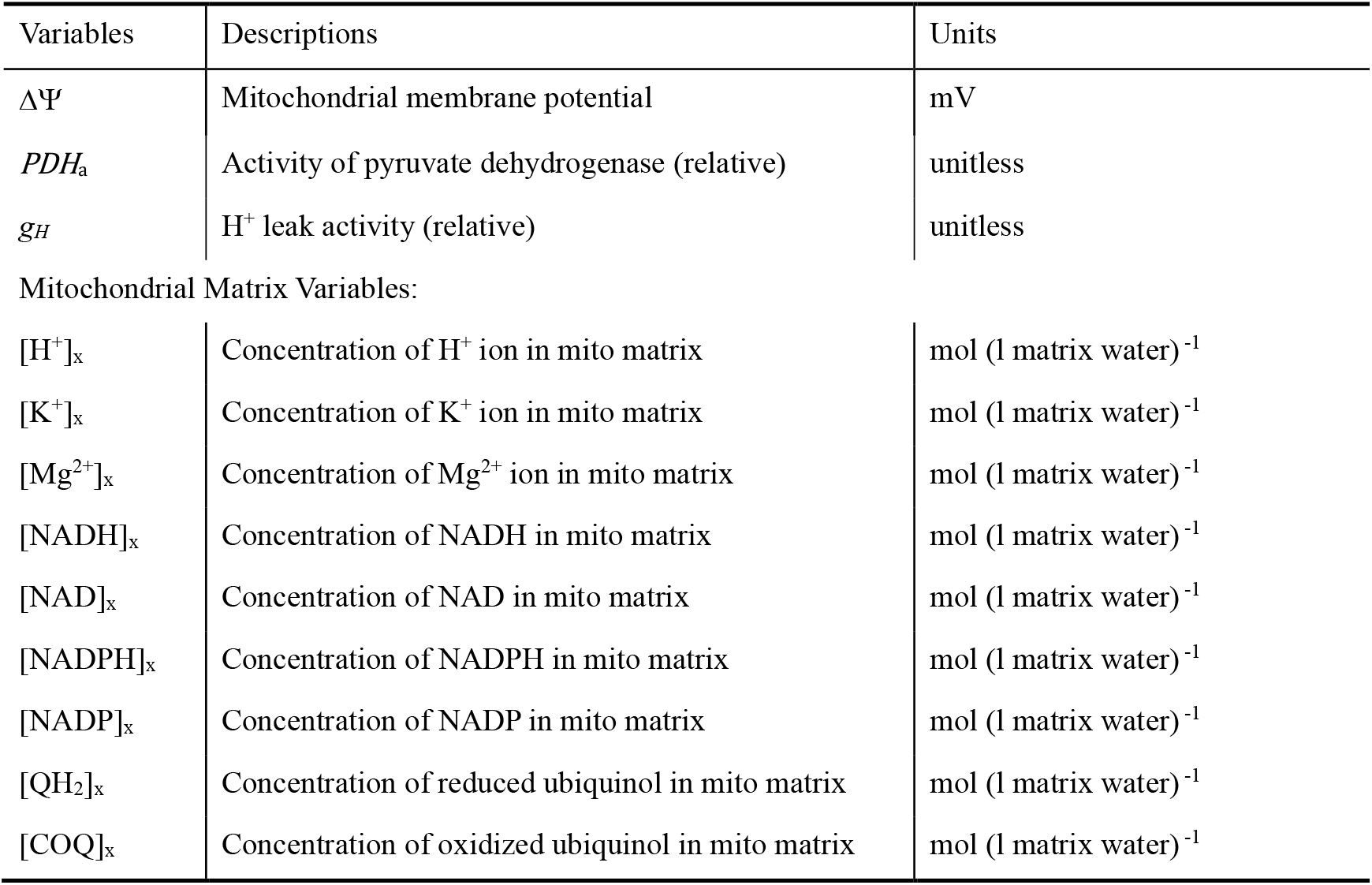
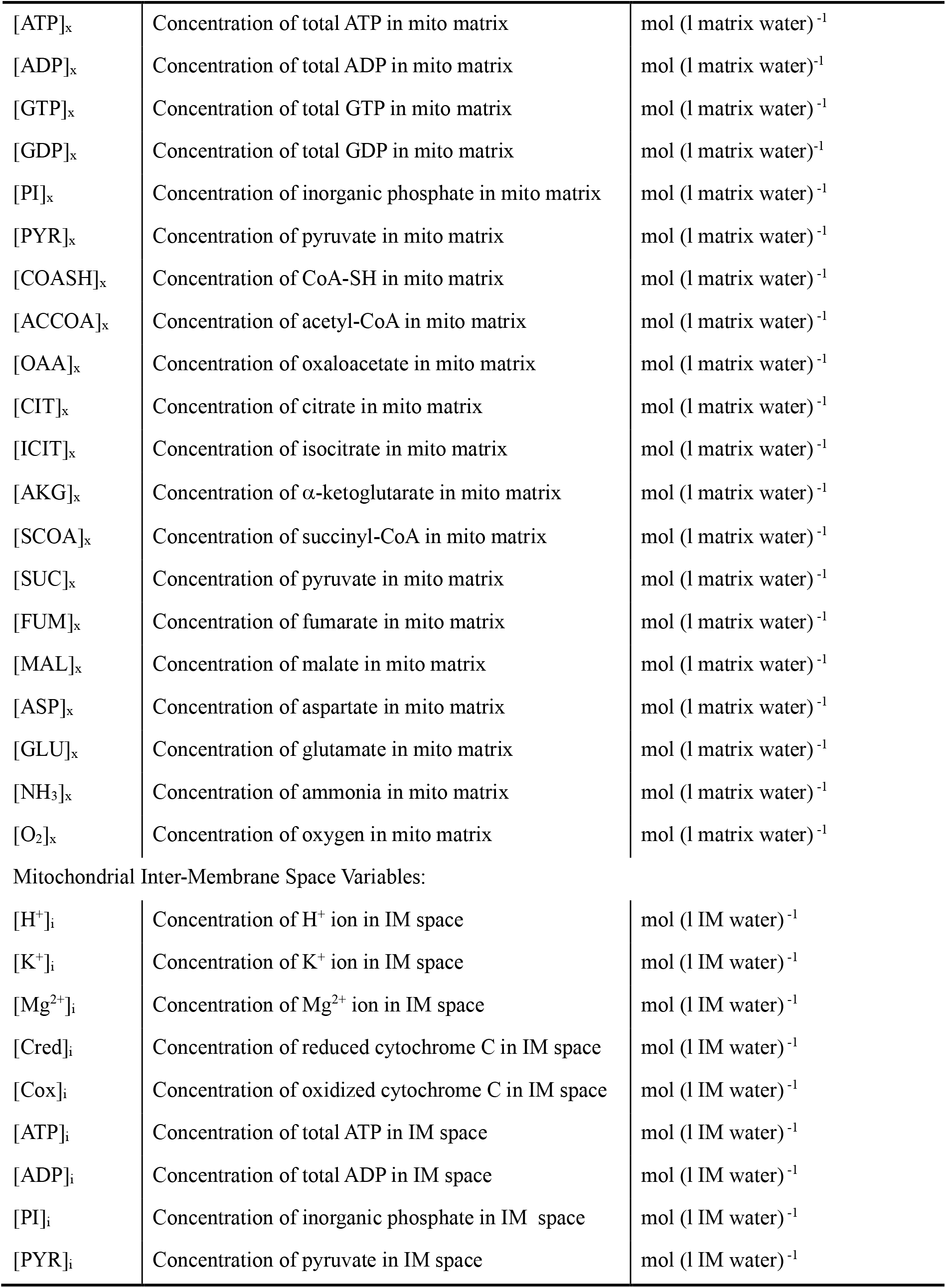
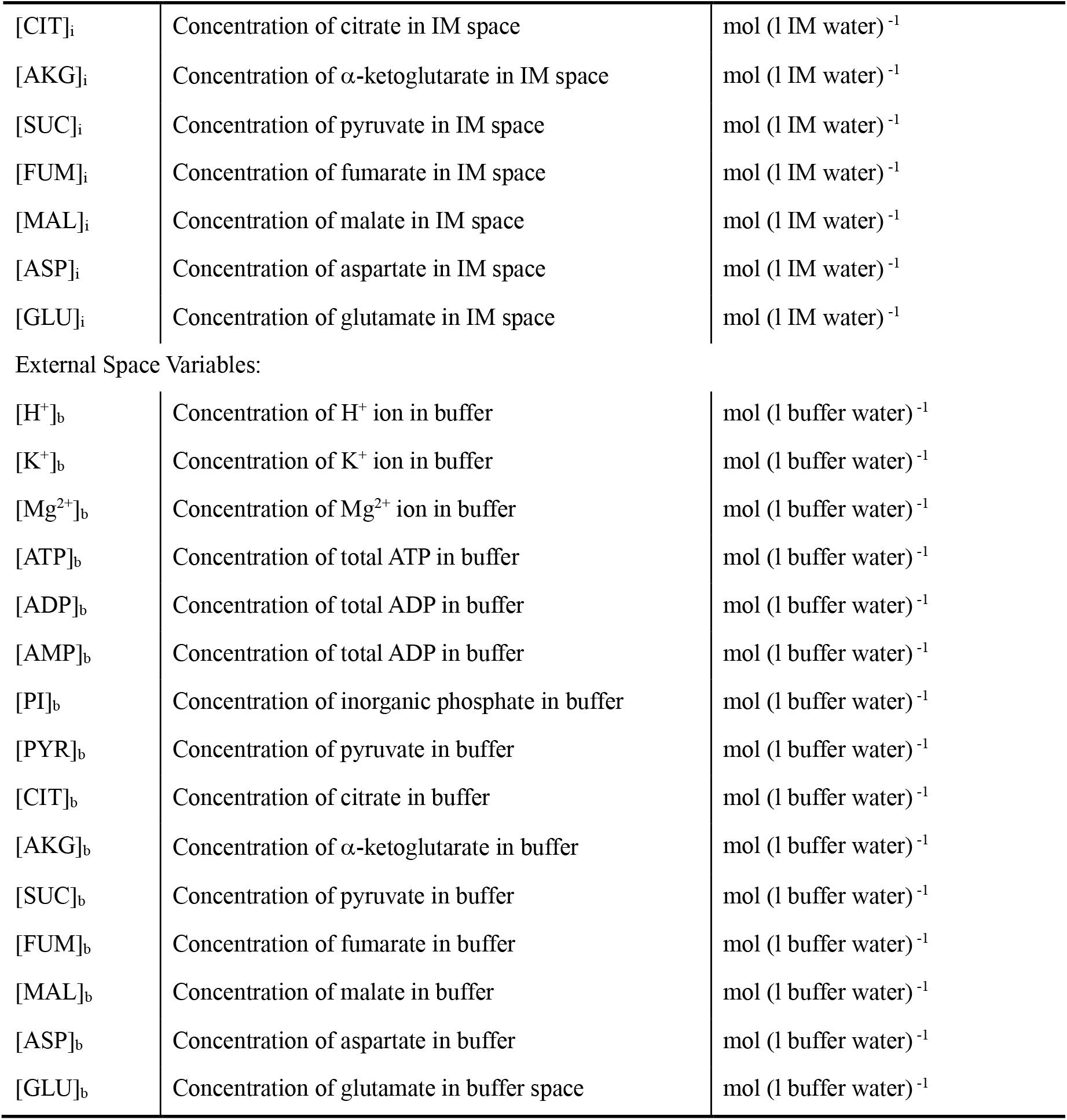
Model Variables.

| Variables | Descriptions | Units |
| --- | --- | --- |
| $\Delta\Psi$ | Mitochondrial membrane potential | mV |
| $PDH_a$ | Activity of pyruvate dehydrogenase (relative) | unitless |
| $g_H$ | H<sup>+</sup> leak activity (relative) | unitless |
| Mitochondrial Matrix Variables: |  |  |
| $[H^+]_x$ | Concentration of H<sup>+</sup> ion in mito matrix | mol (l matrix water)<sup>-1</sup> |
| $[K^+]_x$ | Concentration of K<sup>+</sup> ion in mito matrix | mol (l matrix water)<sup>-1</sup> |
| $[Mg^{2+}]_x$ | Concentration of Mg<sup>2+</sup> ion in mito matrix | mol (l matrix water)<sup>-1</sup> |
| $[NADH]_x$ | Concentration of NADH in mito matrix | mol (l matrix water)<sup>-1</sup> |
| $[NAD]_x$ | Concentration of NAD in mito matrix | mol (l matrix water)<sup>-1</sup> |
| $[NADPH]_x$ | Concentration of NADPH in mito matrix | mol (l matrix water)<sup>-1</sup> |
| $[NADP]_x$ | Concentration of NADP in mito matrix | mol (l matrix water)<sup>-1</sup> |
| $[QH_2]_x$ | Concentration of reduced ubiquinol in mito matrix | mol (l matrix water)<sup>-1</sup> |
| $[COQ]_x$ | Concentration of oxidized ubiquinol in mito matrix | mol (l matrix water)<sup>-1</sup> |
| [ATP]<sub>x</sub> | Concentration of total ATP in mito matrix | mol (l matrix water)<sup>-1</sup> |
| [ADP]<sub>x</sub> | Concentration of total ADP in mito matrix | mol (l matrix water)<sup>-1</sup> |
| [GTP]<sub>x</sub> | Concentration of total GTP in mito matrix | mol (l matrix water)<sup>-1</sup> |
| [GDP]<sub>x</sub> | Concentration of total GDP in mito matrix | mol (l matrix water)<sup>-1</sup> |
| [PI]<sub>x</sub> | Concentration of inorganic phosphate in mito matrix | mol (l matrix water)<sup>-1</sup> |
| [PYR]<sub>x</sub> | Concentration of pyruvate in mito matrix | mol (l matrix water)<sup>-1</sup> |
| [COASH]<sub>x</sub> | Concentration of CoA-SH in mito matrix | mol (l matrix water)<sup>-1</sup> |
| [ACCOA]<sub>x</sub> | Concentration of acetyl-CoA in mito matrix | mol (l matrix water)<sup>-1</sup> |
| [OAA]<sub>x</sub> | Concentration of oxaloacetate in mito matrix | mol (l matrix water)<sup>-1</sup> |
| [CIT]<sub>x</sub> | Concentration of citrate in mito matrix | mol (l matrix water)<sup>-1</sup> |
| [ICIT]<sub>x</sub> | Concentration of isocitrate in mito matrix | mol (l matrix water)<sup>-1</sup> |
| [AKG]<sub>x</sub> | Concentration of α-ketoglutarate in mito matrix | mol (l matrix water)<sup>-1</sup> |
| [SCOA]<sub>x</sub> | Concentration of succinyl-CoA in mito matrix | mol (l matrix water)<sup>-1</sup> |
| [SUC]<sub>x</sub> | Concentration of pyruvate in mito matrix | mol (l matrix water)<sup>-1</sup> |
| [FUM]<sub>x</sub> | Concentration of fumarate in mito matrix | mol (l matrix water)<sup>-1</sup> |
| [MAL]<sub>x</sub> | Concentration of malate in mito matrix | mol (l matrix water)<sup>-1</sup> |
| [ASP]<sub>x</sub> | Concentration of aspartate in mito matrix | mol (l matrix water)<sup>-1</sup> |
| [GLU]<sub>x</sub> | Concentration of glutamate in mito matrix | mol (l matrix water)<sup>-1</sup> |
| [NH<sub>3</sub>]<sub>x</sub> | Concentration of ammonia in mito matrix | mol (l matrix water)<sup>-1</sup> |
| [O<sub>2</sub>]<sub>x</sub> | Concentration of oxygen in mito matrix | mol (l matrix water)<sup>-1</sup> |
| Mitochondrial Inter-Membrane Space Variables: |  |  |
| [H<sup>+</sup>]<sub>i</sub> | Concentration of H<sup>+</sup> ion in IM space | mol (l IM water)<sup>-1</sup> |
| [K<sup>+</sup>]<sub>i</sub> | Concentration of K<sup>+</sup> ion in IM space | mol (l IM water)<sup>-1</sup> |
| [Mg<sup>2+</sup>]<sub>i</sub> | Concentration of Mg<sup>2+</sup> ion in IM space | mol (l IM water)<sup>-1</sup> |
| [Cred]<sub>i</sub> | Concentration of reduced cytochrome C in IM space | mol (l IM water)<sup>-1</sup> |
| [Cox]<sub>i</sub> | Concentration of oxidized cytochrome C in IM space | mol (l IM water)<sup>-1</sup> |
| [ATP]<sub>i</sub> | Concentration of total ATP in IM space | mol (l IM water)<sup>-1</sup> |
| [ADP]<sub>i</sub> | Concentration of total ADP in IM space | mol (l IM water)<sup>-1</sup> |
| [PI]<sub>i</sub> | Concentration of inorganic phosphate in IM space | mol (l IM water)<sup>-1</sup> |
| [PYR]<sub>i</sub> | Concentration of pyruvate in IM space | mol (l IM water)<sup>-1</sup> |
| [CIT]<sub>i</sub> | Concentration of citrate in IM space | mol (l IM water)<sup>-1</sup> |
| [AKG]<sub>i</sub> | Concentration of α-ketoglutarate in IM space | mol (l IM water)<sup>-1</sup> |
| [SUC]<sub>i</sub> | Concentration of pyruvate in IM space | mol (l IM water)<sup>-1</sup> |
| [FUM]<sub>i</sub> | Concentration of fumarate in IM space | mol (l IM water)<sup>-1</sup> |
| [MAL]<sub>i</sub> | Concentration of malate in IM space | mol (l IM water)<sup>-1</sup> |
| [ASP]<sub>i</sub> | Concentration of aspartate in IM space | mol (l IM water)<sup>-1</sup> |
| [GLU]<sub>i</sub> | Concentration of glutamate in IM space | mol (l IM water)<sup>-1</sup> |
| External Space Variables: |  |  |
| [H<sup>+</sup>]<sub>b</sub> | Concentration of H<sup>+</sup> ion in buffer | mol (l buffer water)<sup>-1</sup> |
| [K<sup>+</sup>]<sub>b</sub> | Concentration of K<sup>+</sup> ion in buffer | mol (l buffer water)<sup>-1</sup> |
| [Mg<sup>2+</sup>]<sub>b</sub> | Concentration of Mg<sup>2+</sup> ion in buffer | mol (l buffer water)<sup>-1</sup> |
| [ATP]<sub>b</sub> | Concentration of total ATP in buffer | mol (l buffer water)<sup>-1</sup> |
| [ADP]<sub>b</sub> | Concentration of total ADP in buffer | mol (l buffer water)<sup>-1</sup> |
| [AMP]<sub>b</sub> | Concentration of total ADP in buffer | mol (l buffer water)<sup>-1</sup> |
| [PI]<sub>b</sub> | Concentration of inorganic phosphate in buffer | mol (l buffer water)<sup>-1</sup> |
| [PYR]<sub>b</sub> | Concentration of pyruvate in buffer | mol (l buffer water)<sup>-1</sup> |
| [CIT]<sub>b</sub> | Concentration of citrate in buffer | mol (l buffer water)<sup>-1</sup> |
| [AKG]<sub>b</sub> | Concentration of α-ketoglutarate in buffer | mol (l buffer water)<sup>-1</sup> |
| [SUC]<sub>b</sub> | Concentration of pyruvate in buffer | mol (l buffer water)<sup>-1</sup> |
| [FUM]<sub>b</sub> | Concentration of fumarate in buffer | mol (l buffer water)<sup>-1</sup> |
| [MAL]<sub>b</sub> | Concentration of malate in buffer | mol (l buffer water)<sup>-1</sup> |
| [ASP]<sub>b</sub> | Concentration of aspartate in buffer | mol (l buffer water)<sup>-1</sup> |
| [GLU]<sub>b</sub> | Concentration of glutamate in buffer space | mol (l buffer water)<sup>-1</sup> |

**Table A2:**
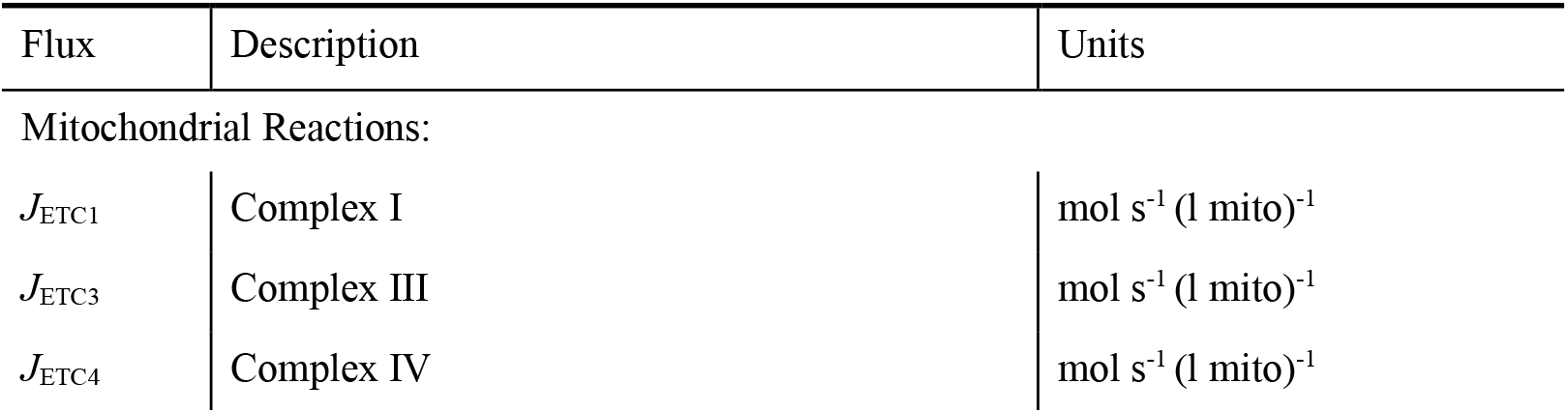
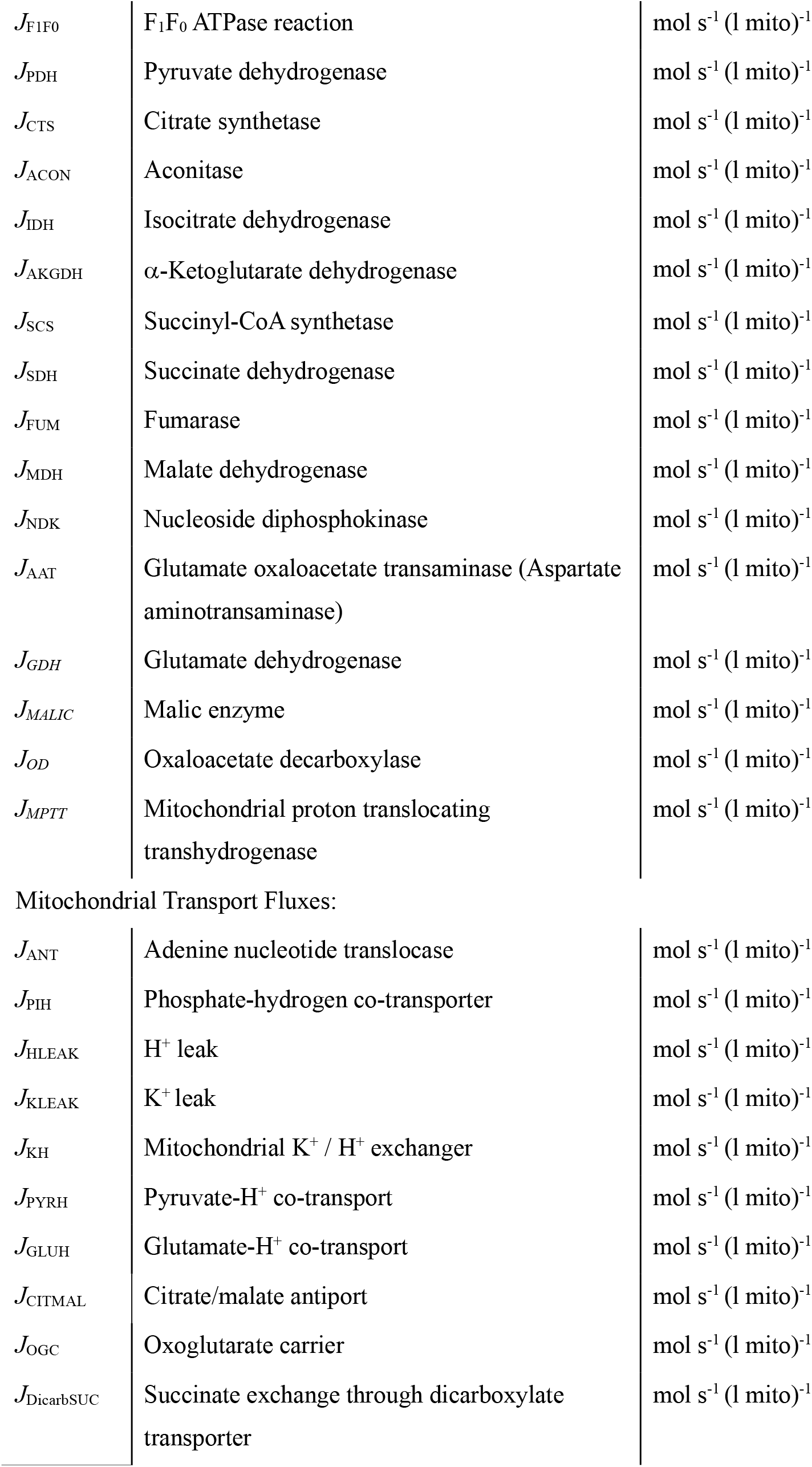
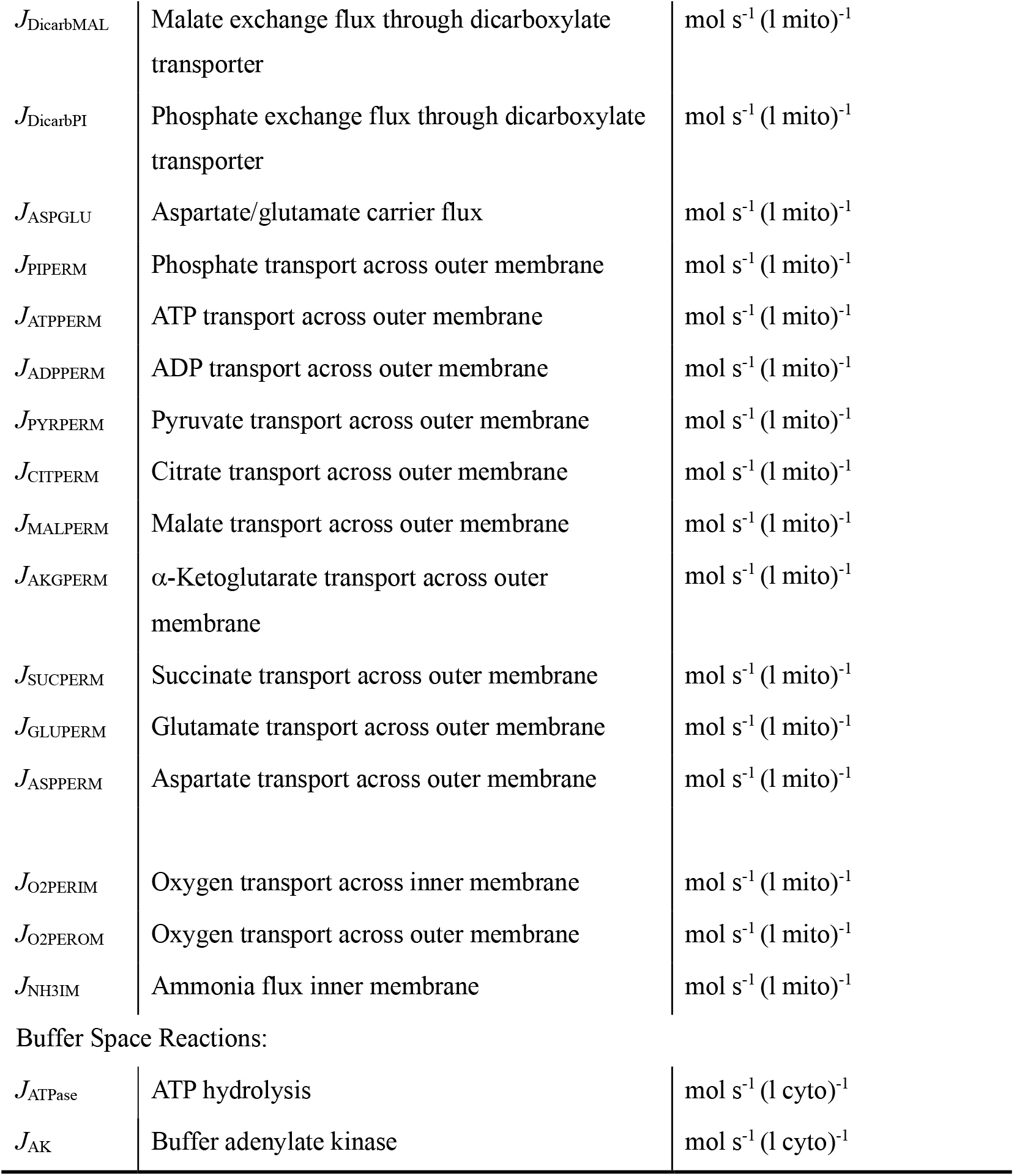
Reaction and Transport Fluxes.

**Table A3:**
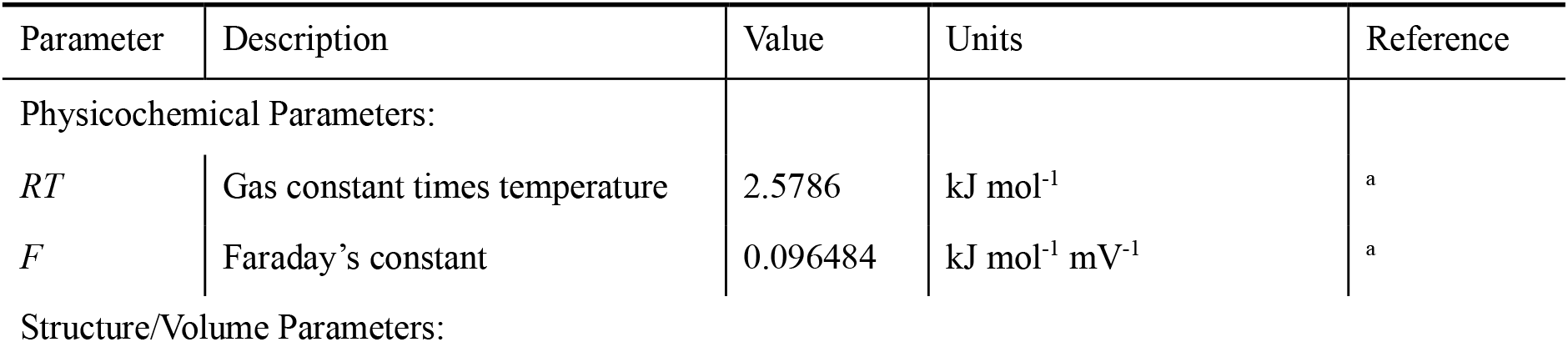
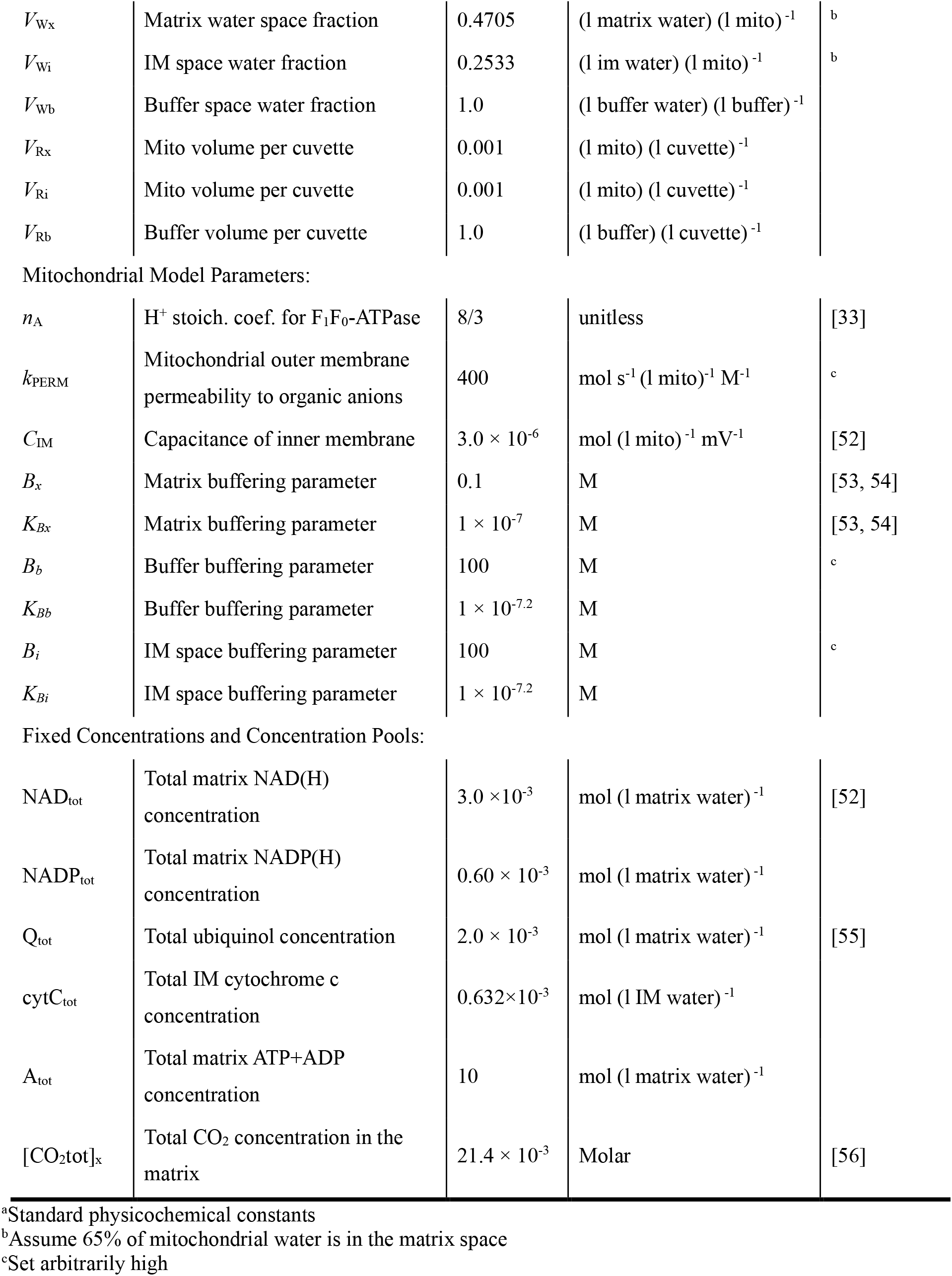
Fixed Model Parameters.

### C. Governing Equations

The model is mathematically described by using the differential equations listed in this appendix. The oxidative phosphorylation component of the model is derived from previously published work [33]. The details behind the TCA cycle enzyme kinetic schemes are provided in Appendix C. Here the subscripts “x”, “i”, and “b” on variable names denote matrix, intermembrane, and buffer (extra-mitochondrial) spaces, respectively. For example [ATP]_x_ denotes matrix ATP concentration while [ATP]_b_ denotes ATP concentration in the buffer space for an isolated mitochondria experiment.

The differential equations are grouped into equations for membrane potential, mitochondrial matrix variables, intermembrane space variables, and external space variables. The external space variables used depend on the type of simulation considered (ex vivo isolated mitochondrial or the in vivo system.)

#### Mitochondrial Inner Membrane Electrical Potential

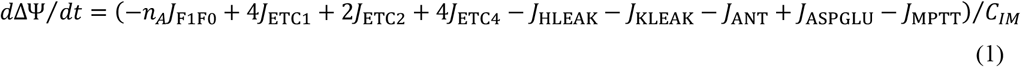

#### Mitochondrial Matrix

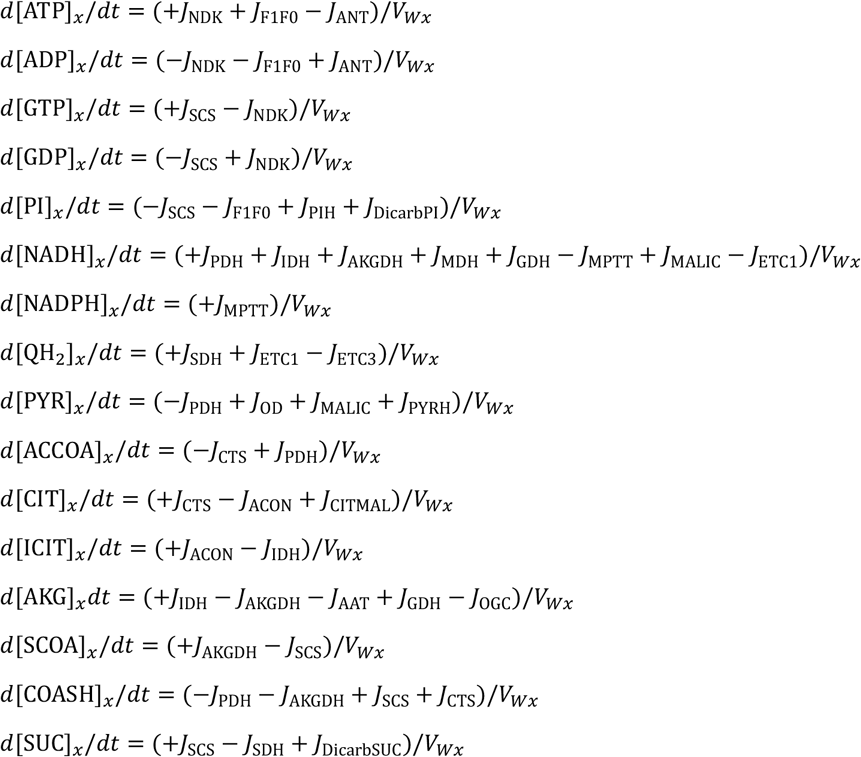

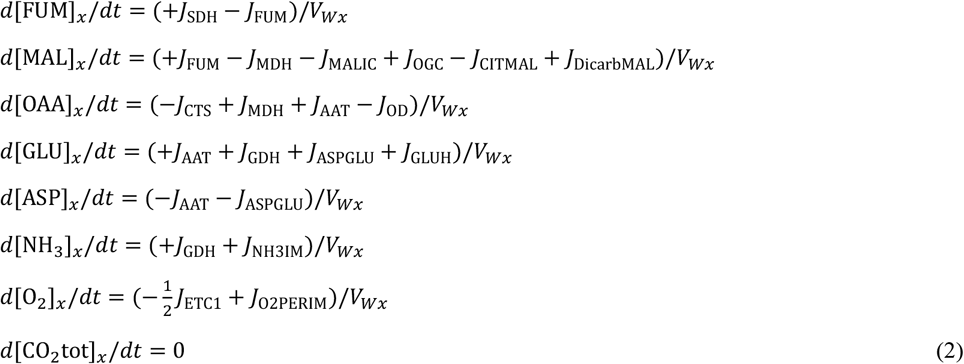

#### Mitochondrial Inter-Membrane Space

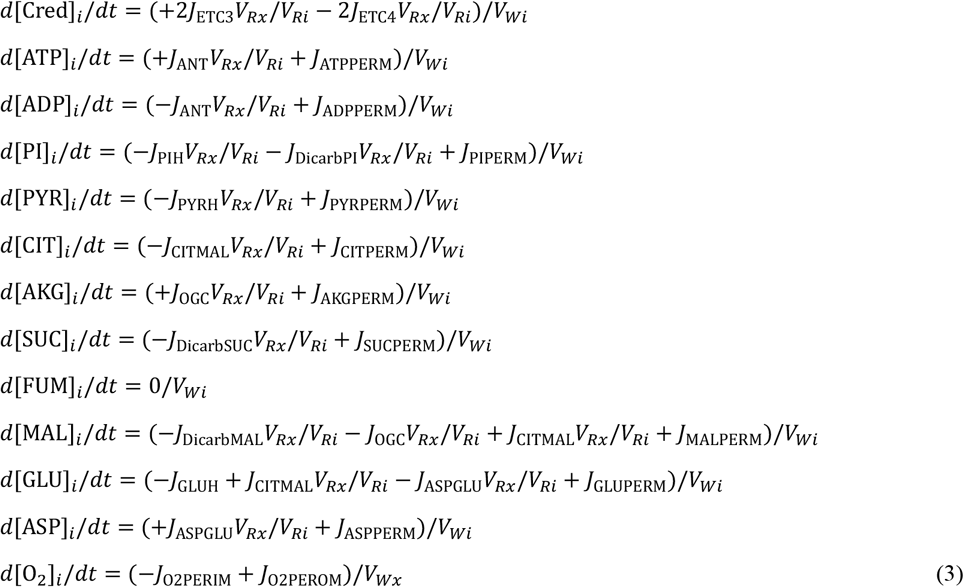

#### External Space

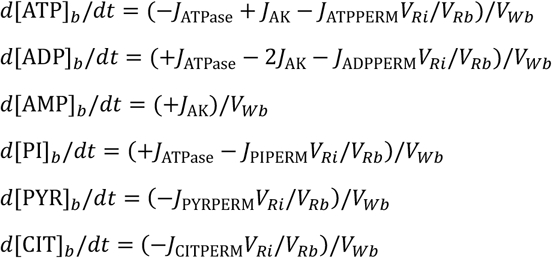

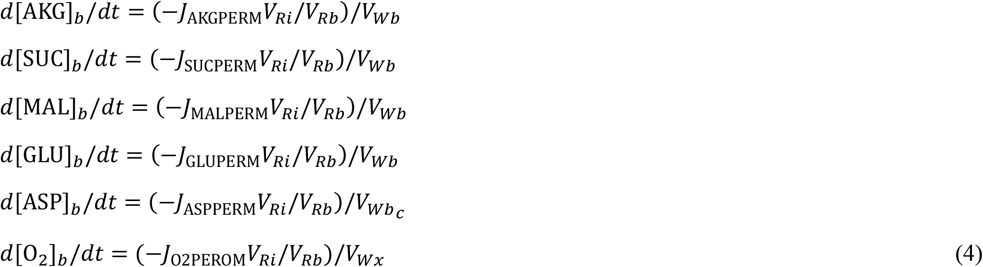

Assuming constant total concentrations NAD_tot_, Q_tot_, cytC_tot_, and A_tot_ for nicotinamide nucleotides, ubiquinol, and cytochrome c, we compute concentrations of the following reactants as:

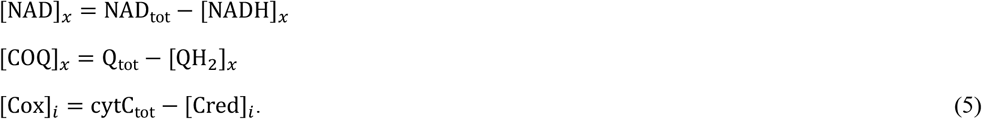

#### Pyruvate dehydrogenase activity

Pyruvate dehydrogenase activity, represented by *PDH*_a_(*t*), is governed by a phosphorylation dephosphorylation cycle, where *PDH*_a_(*t*) = 1 at maximal activity, and *PDH*_a_(*t*) = 0 represents the fully phosphorylated inactive state. Activation of PDH is assumed inhibited by NADH, while inactivation is stimulated by NADH and requires ATP:

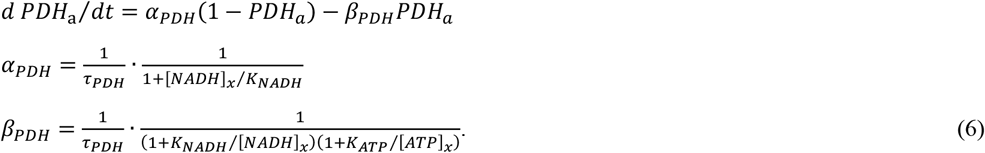

This simplified model invokes three adjustable parameters, *τ*_PDH_ = 87.78 seconds, *K*_NADH_ = 0.484 × 10^−3^ M, *K*_ATP_ = 0.964 × 10^−3^ M, which are estimated based on matching the oxygen consumption and NAD(P)H time courses for experiments using pyruvate as a substrate.

#### H^+^ leak activity

Proton ion leak activity, represented by *g*_H_(*t*), is assumed to be activated by high levels of reactive oxygen species production. To account for this phenomenon, the steady-state leak conductance is taken to be proportional to the reduction state of the ubiquinol pool:

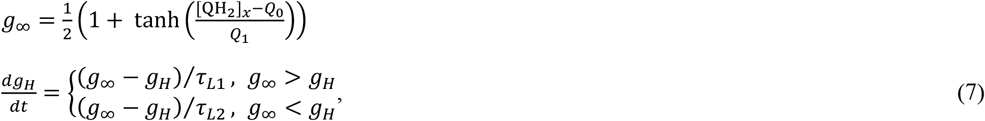

where increased QH_2_ level (associated with increased ROS production) leads to increased activity of UCP3, and increased proton leak conductance. The parameters *Q*_0_ = *Q*_1_ = 0.25*Q*_*tot*_ are set such that the transporter remains largely closed under conditions of respiration fueled by complex-I substrates and activates under succinate-fueled respiration. The time constant *τ*_*L*1_ = 5 seconds is arbitrarily set to achieve rapid activation on the timescales observed. The time constant *τ*_*L*2_ = 225 seconds, associated with deactivation of the transporter, is estimated based on fitting model simulations to data from succinate-fueled conditions.

#### Governing equations for pH, Mg^2+^, and K^+^

Governing equations for [H+], [Mg^2+^], and [K^+^] in each compartment follow from [14-16, 23]. In each compartment, the kinetics of pH, [Mg^2+^], and [K^+^] are governed by cation binding and unbinding as well as the consumption and generation of protons via chemical reactions. For a full derivation and details see Wu et al. [15] and Vinnakota et al. [23].

The time derivatives of [H^+^], [Mg^2+^], and [K^+^] are given by

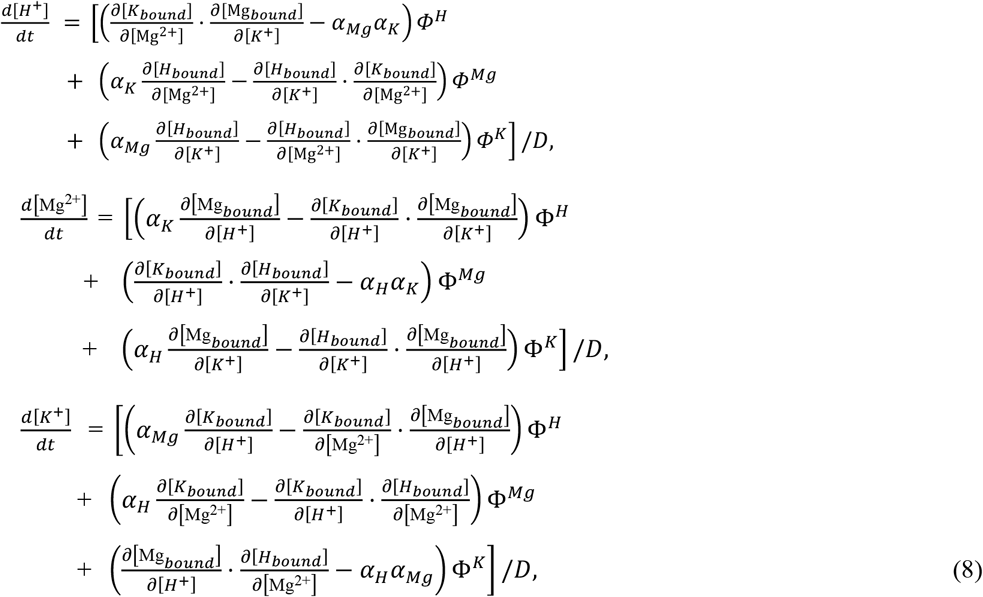

where

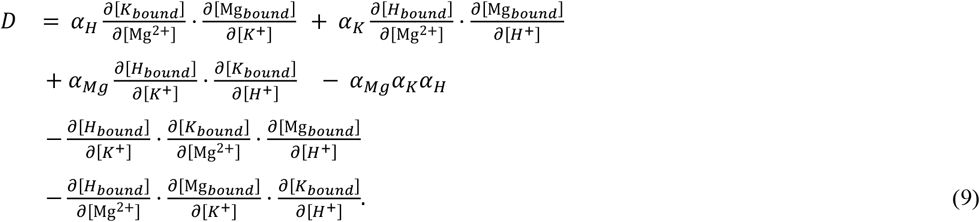

The variables in Equations (8) and (9) are defined

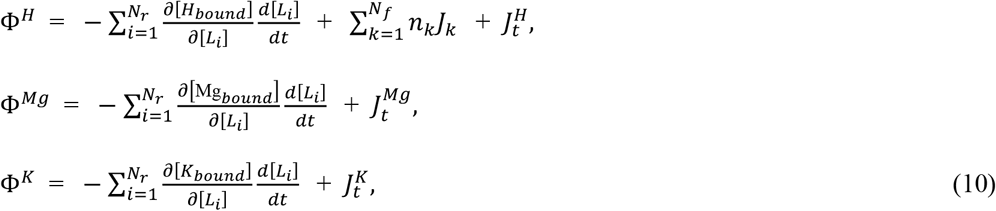

where *N*_*r*_ is the number of reactants, *N*_*f*_ is the number of reactions, *n*_*k*_ is the stoichiometric coefficient of H^+^ for *k*^*th*^ reaction, *J*_*k*_ is the flux of *k*^*th*^ reaction, 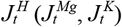 is the transport flux of [H^+^] ([Mg^2+^], [K^+^]) into the compartment. The concentrations [H_bound_], [K_bound_], and [Mg_bound_] are the concentrations of H^+^, Mg^2+^, K^+^ reversibly bound to organic cations on the system. For example,

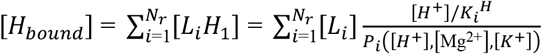

where [L_*i*_] is the total concentration for reactant *i*, 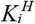 and *P*_*i*_ are the proton binding constant and binding polynomial for the reactant. The binding polynomials are calculated as

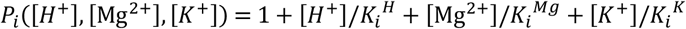

where 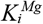 and 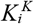 are the binding constants for Mg^2+^ and K^+^. The partial derivatives are

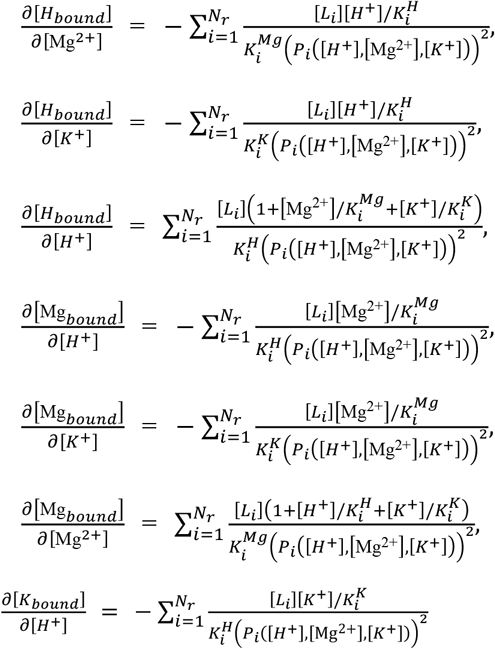

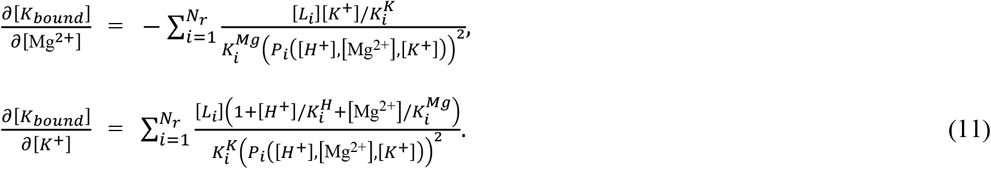

Finally, the buffering terms are:

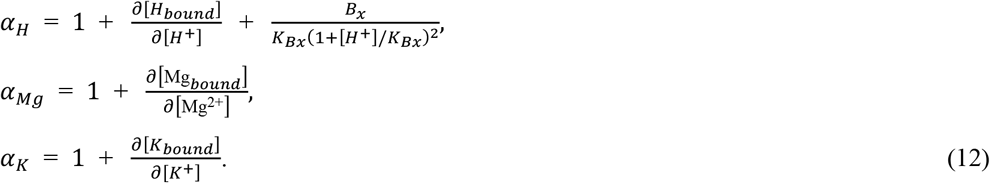

### D. NADH Binding

The model represents NADH binding to many mitochondrial enzymes through a simplified first-order binding process that captures the combined effects of multiple underlying interactions:

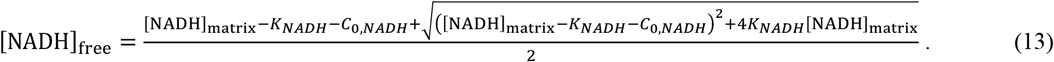

The model assumes an overall binding capacity of *C*_0,*NADH*_ = 1 mM. The effective dissociation constant is estimated *K*_*NADH*_ = 0.157 × 10^−3^ M based on fitting model simulations to data.

### E. Flux Expressions

#### Complex I and Complex III

Flux expressions for the complex I and III components of the respiratory chain are adopted from Bazil et al. [34, 35], and implemented as in the model of Bazil et al. [33]. Adjustable parameters representing the complex I and III content in mitochondria, Etot_C1_ = 0.229 × 10^−3^ mol (l mito)^-1^ and Etot_C3_ = 0.639 × 10^−3^ mol (l mito)^-1^ are identified based on fitting model simulations to data.

#### Complex IV

The complex IV flux is modeled using the phenomenological expression determined by Bazil et al. [33]:

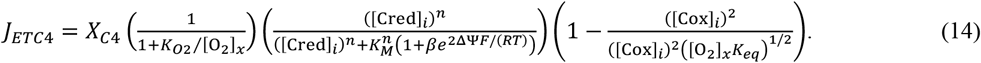

Parameter values are *K*_*O*2_ = 1 × 10^−6^ M, *K*_*M*_ = 162 × 10^−6^ M, *n* = 2, *β* = 6.6 × 10^−6^, and *X*_*C*4_ = 3.75 × 10^−3^ mol s^-1^ (liter mito volume)^-1^. The parameter *X*_*C*4_ is identified based on fitting model simulations to data.

#### F_1_F_0_ ATPase reaction

Previous models have assumed a simple mass-action formulation for the F_1_F_0_ ATPase reaction. Here we assume a simple reversible random bi-bi model:

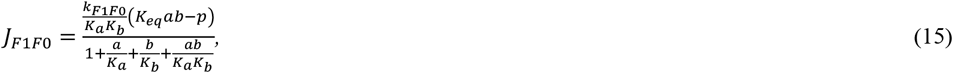

where *a* = [ADP]_x_, *b* = [Pi]_x_, and *p* = [ATP]_x_. Parameters *K*_*a*_ and *K*_*b*_ are arbitrary set to *Ka* = 1 × 10^−3^ M *K*_*b*_ = 2 × 10^−3^ M and The parameter *k*_*F*1*F*0_ = 0.38883 × 10^−3^ mol s^-1^ M^-1^ (liter mito volume)^-1^ is identified based on fitting model simulations to data.

#### Pyruvate dehydrogenase

The flux expression follows from Wu et al. [15], modified to incorporate the time-varying activity governed by Equation (6):

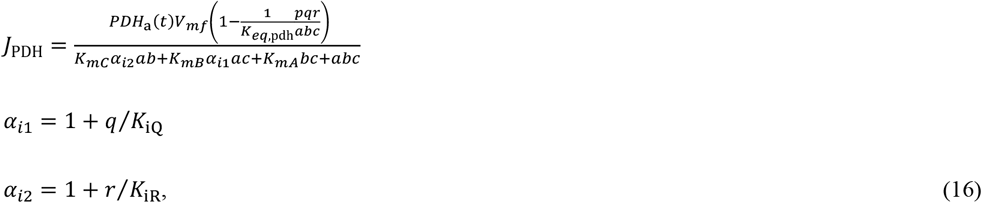

where *a* = [PYR]_x_, *b* = [COASH]_x_, *c* = [NAD]_free_, *p* = [CO_2_tot]_x_, *q* = [ACCOA]_x_, *r* = [NADH]_free_, and parameter values are *V*_mf_ = 3.455 × 10^−3^ mol s^-1^ (liter mito volume)^-1^, *K*_mC_ = 0.91132 × 10^−3^ M, *K*_mA_ = 38.3 × 10^−6^ M, *K*_mB_ = 9.9 × 10^−6^ M, *K*_iQ_ = 40.2 × 10^−6^ M, and *K*_iR_ = 0.13441 × 10^−3^ M.

The parameters *V*_mf_, *K*_mC_, and *K*_iR_ are identified based on fitting model simulations to data.

#### Citrate synthase

The flux expression follows from Wu et al. [26]:

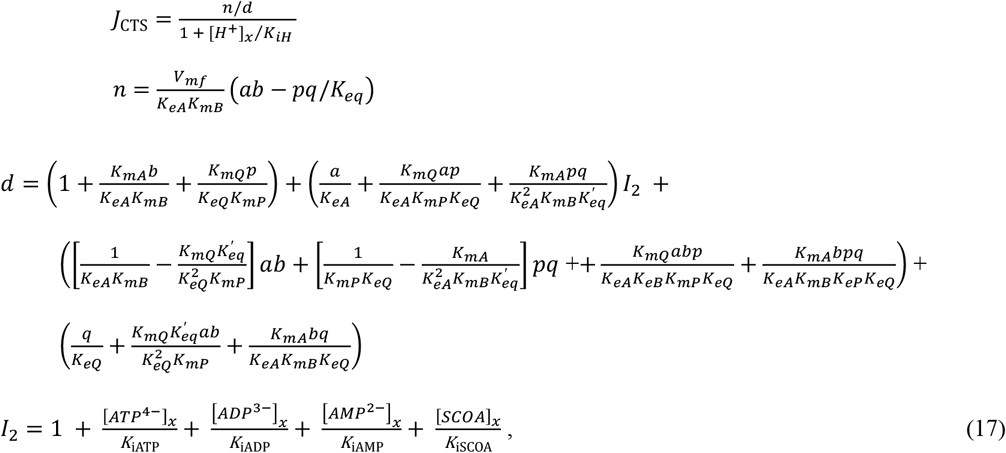

where *a* = [OAA]_x_, *b* = [ACCOA]_x_, *p* = [COASH]_x_, *q* = [CIT]_x_, and parameter values are *V*_mf_ = 0.1676 mol s^-1^ (liter mito volume)^-1^, *K*_mA_ = 5.7628 × 10^−6^ M, *K*_mB_ = 4.5728 × 10^−6^ M, *K*_mP_ = 0.1788 × 10^−6^ M, *K*_MQ_ = 2.24 × 10^−3^ M, *K*_eA_ = 3.0898 × 10^−6^ M, *K*_eB_ = 0.283 M, *K*_eQ_ = 0.4475 × 10^−3^ M, *K*_iATP_ = 39.898 × 10^−6^ M, *K*_iADP_ = 141.79 × 10^−6^ M, *K*_iAMP_ = 1.0533 × 10^−3^ M, *K*_iSCOA_ = 8.9669 × 10^−6^ M, *K*_iH_ = 5.5 × 10^−8^ M, and

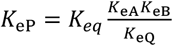

The parameter *V*_mf_ is identified based on fitting model simulations to data.

#### Aconitase

The flux expression follows from Wu et al. [15]:

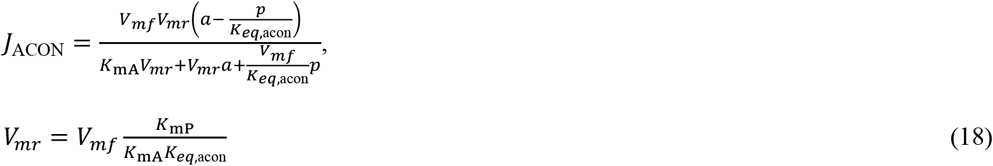

where *a* = [CIT]_x_ and *p* = [ICIT]_x_, and parameter values are *K*_mA_ = 1161 × 10^−6^ M, *K*_mP_ = 434 × 10^−6^ M. The parameter *V*_mf_ = 0.28613 mol s^-1^ (liter mito volume)^-1^ is identified based on fitting model simulations to data.

#### Isocitrate dehydrogenase

The flux expression follows from Qi et al. [27]:

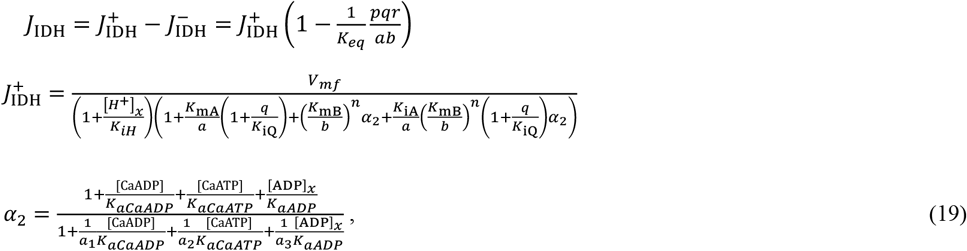

where *a* = [NAD]_x_, *b* = [ICIT]_x_, *p* = [AKG]_x_, *q* = [NADH]_free_, and *r* = [CO_2_tot]_x_. The concentrations [CaADP] and [CaATP] represent Ca^2+^-bound ADP and ATP, computed:

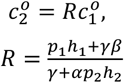

where *K*_*CaATP*_ = 0.138 × 10^−3^ M and *K*_*CaATP*_ = 1.38 × 10^−3^ M are dissociation constants. Parameter values are *V*_*mf*_ = 2.207 × 10^−3^ mol s^-1^ (liter mito volume)^-1^, *K*_*iH*_ = 0.11 × 10^−6^ M, *K*_*mA*_ = 503.3 × 10^−6^ M, *K*_*iB*_ 148.9 × 10^−6^ M, *K*_*iA*_ = 77.6 × 10^−6^ M, *K*_*aCaADP*_ = 13.3 × 10^−6^ M, *K*_*aCaATP*_ = 288.6 × 10^−6^ M, *K*_*aADP*_ = 61.3 × 10^−3^ M, *K*_*iQ*_ = 4.75 × 10^−6^ M, *a*_1_ = 0.0012, *a*_2_= 0.0097, *a*_1_= 0.0004, and *n* = 3.

The parameter *V*_*m*f_is identified based on fitting model simulations to data.

#### α-Ketoglutarate dehydrogenase

The flux expression follows from Qi et al. [28]:

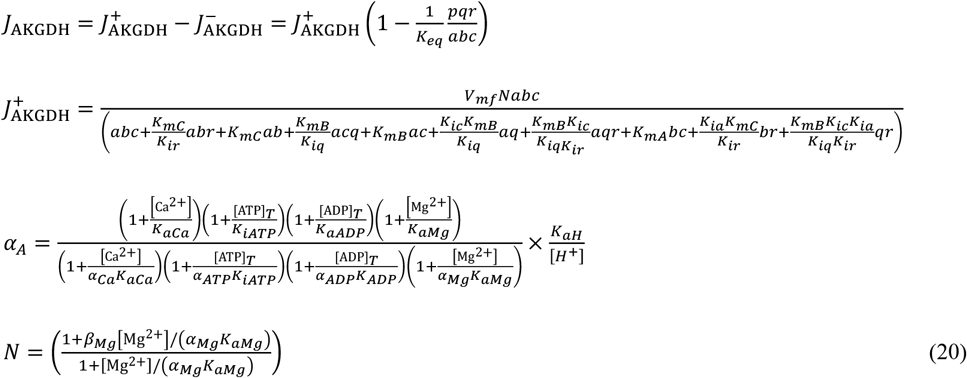

where *a* = [AKG]_x_, *b* = [COASH]_x_, *c* = [NAD]_x_, *p* = [CO_2_tot]_x_, *q* = [SCOA]_x_, and *r* = [NADH]_free_. Parameter values are *V*_*mf*_= 2.8437 × 10^−3^ mol s^-1^ (liter mito volume)^-1^, *K*_*mA*_= 0.273 × 10^−3^ M, *K*_*iB*_= 6.96 × 10^−6^ M, *K*_*mC*_= 98.6 × 10^−6^ M, *K*_*ia*_= 75.9 × 10^−3^ M, *K*_*ir*_= 2.4 × 10^−6^ M, *K*_*ic*_= 0.112 × 10^−3^ M, *K*_*iq*_= 0.218 × 10^−3^ M, *K*_*aH*_= 7.763 × 10^−7^ M, *K*_*aCa*_= 0.893 × 10^−6^ M, *K*_*iATP*_= 0.106 × 10^−3^ M, *K*_*iATP*_= 0.3055 × 10^−3^ M, *K*_*aMg*_= 19.49 × 10^−6^ M, *α*_*Ca*_= 0.262, *α*_*ATP*_= 6.674, *α*_*ADP*_= 0.173, *α*_*Mg*_= 1, and *β*_*Mg*_= 4.222.

The parameter *V*_*mf*_is identified based on fitting model simulations to data.

#### Succinyl-CoA synthetase

The flux expression follows from Li et al. [29]:

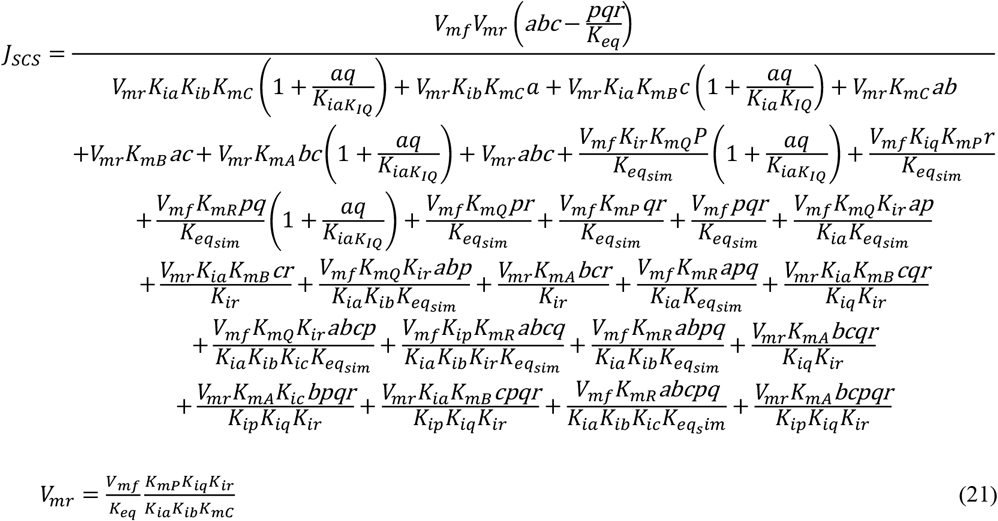

where for the Li et al. model, *a, b, b, p, q*, and *r* represent the activities of the species involved in the reaction, estimated as a function of temperature *T* and ionic strength *I*:

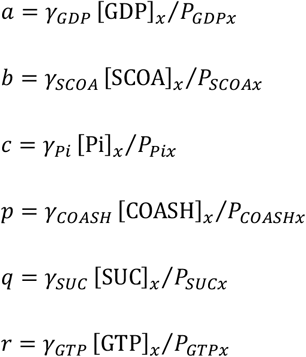

where the activity coefficient *γ* is e The activity coefficient *γ*_*i*_for a given species is be estimated from

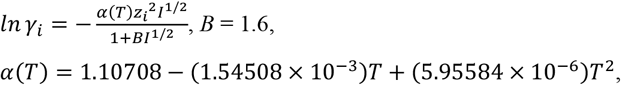

and *z*_*i*_is the valence of species *i*. Parameter values are *V*_*mf*_= 6.2297 × 10^−3^ mol s^-1^ (liter mito volume)^-1^, *K*_*iQ*_= 1.484 × 10^−4^ M, *K*_*mA*_= 2.862 × 10^−9^ M, *K*_*mB*_= 7.807 × 10^−6^ M, *K*_*mC*_= 3.821 × 10^−4^ M, *K*_*mP*_= 5.109 × 10^−7^ M, *K*_*mQ*_= 6.510 × 10^−5^ M, *K*_*mR*_= 5.567 × 10^−10^ M, *K*_*ia*_= 2.707 × 10^−8^ M, *K*_*ib*_= 8.699 × 10^−6^ M, *K*_*ic*_= 1.777 × 10^−3^ M, *K*_*ip*_= 2.871 × 10^−6^ M, *K*_*iq*_= 1.759 × 10^−3^ M, and *K*_*ir*_= 7.115 × 10^−10^ M.

The parameter *V*_*mf*_is identified based on fitting model simulations to data.

#### Succinate dehydrogenase

The flux expression follows from Manhas et al. [30]. The flux expression for *J*_SDH_, computed as a function of reactants [QH_2_]_x_, [COQ]_x_, [FUM]_x_, and [SUC]_x_is given by Eq. S93 in the appendix to Manhas et al. [30]. In addition to substrate concentration, the flux depends on matrix pH, as well as concentrations of inhibitors malonate and oxaloacetate.

The maximal flux of succinate dehydrogenase *V*_*mf*_= 0.093131 × 10^−3^ mol s^-1^ (liter mito volume)^-1^ i s identified based on fitting model simulations to data.

#### Fumarase

The flux expression follows from a simplified version of the model of Mescam et al. [32]:

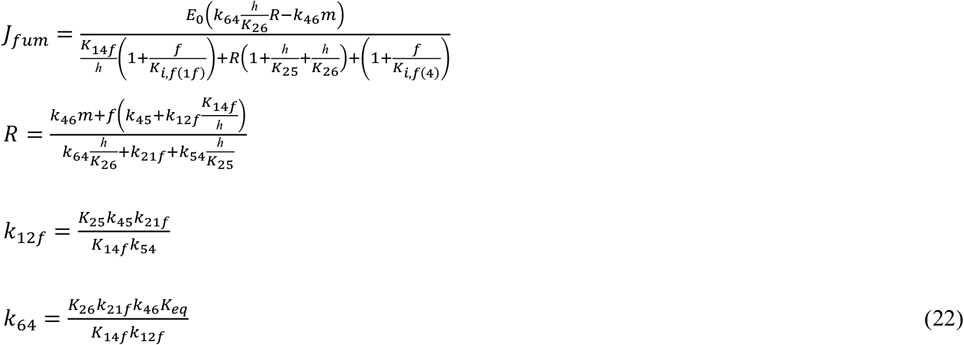

where for the Mescam et al. model, *m* and *f* represent the activities of the species involved in the reaction, estimated as a function of temperature *T* and ionic strength *I*:

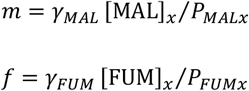

where the activity coefficient *γ* is e The activity coefficient *γ*_*i*_for a given species is be estimated from

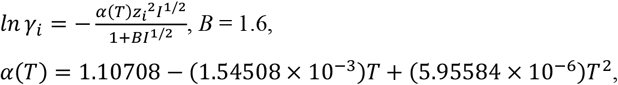

and *z*_*i*_is the valence of species *i*. Parameter values are *E*_0_= 0.15859 × 10^−3^ mol (l mito)^-1^, *k*_46_= 1.91×10^6^ M^-1^·s^-1^, *k*_45_= 4.46×10^5^ M^-1^·s^-1^, *k*_54_= 54.12 s^-1^, *k*_21f_= 5.82×10^4^ s^-1^, *K*_26_= 8.28×10^-8^ M, *K*_25_= 1.81×10^-10^ M, *K*_14f_= 7.69×10^-9^ M.

The parameter *E*_0_is identified based on fitting model simulations to data.

#### Malate Dehydrogenase

The flux expression follows from the model of Dasika et al. [31]:

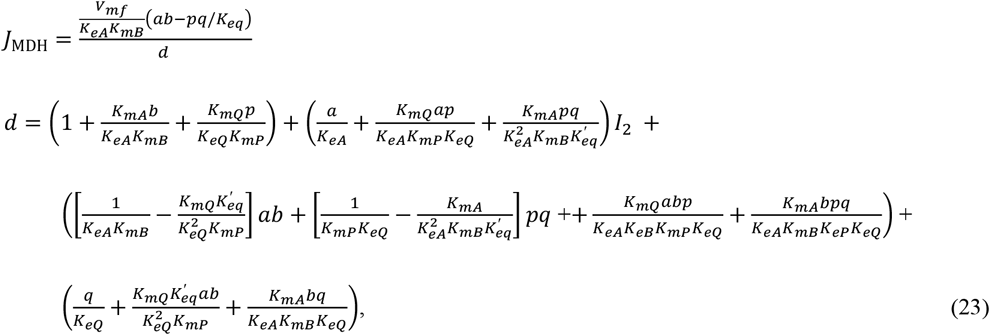

where *a* = [NAD]_x_, *b* = [MAL]_x_, *p* = [OAA]_x_, and *q* = [NADH]_free_. The kinetic constants in Equation (19) are computed as functions of ionic conditions and matrix pH, as described in Dasika et al. [31]. At matrix pH of 7.4, *K*_*mA*_= 7.32 × 10^−5^ M, *K*_*mB*_= 1.69 × 10^−4^ M, *K*_*mP*_= 2.10 × 10^−5^ M, *K*_*mQ*_= 1.06 × 10^−4^ M, *K*_*eA*_= 0.0243, *K*_*eB*_= 4.45 × 10^−4^, *K*_*eP*_= 1.72 × 10^−5^, and *K*_*eQ*_= 1.48 × 10^−5^.

The parameter *V*_*mf*_= 0.051969 mol s^-1^ (liter mito volume)^-1^ is identified based on fitting model simulations to data.

#### Nucleoside diphosphokinase

The nucleoside diphosphokinase model is adopted from Wu et al. [15]:

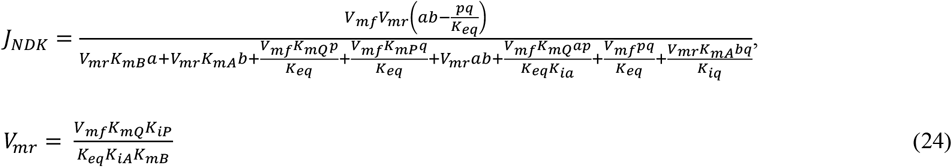

where *K*_*mA*_= 111 × 10^−6^ M, *K*_*mB*_= 100 × 10^−6^ M, *K*_*mP*_= 260 × 10^−6^ M, *K*_*mQ*_= 278 × 10^−6^ M, *K*_*iA*_= 170 × 10^−6^ M, *K*_*iP*_= 146.6 × 10^−6^ M, and *K*_*iQ*_= 156.5 × 10^−6^ M.

The parameter *V*_*mf*_= 0.3857 mol s^-1^ (liter mito volume)^-1^ is identified based on fitting model simulations to data.

#### Glutamate oxaloacetate transaminase

The model for glutamate oxaloacetate transaminase (aspartate amino transferase) is adopted from Saito et al. [18]:

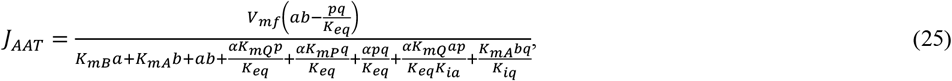

where *a* = [ASP]_x_, *b* =[AKG]_x_, *p* = [OAA]_x_, and *q* = [GLU]_x_. Parameter values are *V*_*mf*_= 0.018126 mol s^-1^ (liter mito volume)^-1^, *α* = 0.517, *K*_*mA*_= 1.58 × 10^−3^ M, *K*_*mB*_= 0.149 × 10^−3^ M, *K*_*mP*_= 0.0399 × 10^−3^ M, *K*_*mQ*_= 2.5 × 10^−3^ M, *K*_*ia*_= 2.0 × 10^−3^ M, *K*_*iq*_= 1.83 × 3 M.

The parameter *V*_*mf*_is identified based on fitting model simulations to data.

#### Glutamate dehydrogenase

The glutamate dehydrogenase model is obtained from Mulukutla et al. [57], based on a random bi-bi mechanism:

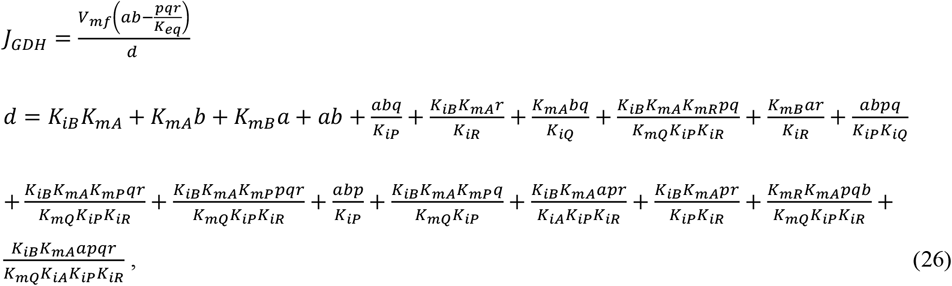

where *a* = [GLU]_x_, *b* =[NAD]_x_, *p* = [AKG]_x_, *r* = [NADH]_free_, and *q* = [NH_2_]_x_. Parameter values are *K*_*mA*_= 3.5 × 10^−3^ M, *K*_*mB*_= 80 × 10^−6^ M, *K*_*mP*_= 1.1 × 10^−3^ M, *K*_*mQ*_= 6.0 × 10^−3^ M, *K*_*mR*_= 40 × 10^−6^ M, *K*_*iA*_= 3.5 × 10^−3^ M, *K*_*iB*_= 1 × 10^−3^ M, *K*_*iP*_= 0.25 × 10^−3^ M, *K*_*iQ*_= 6.0 × 10^−3^ M, *K*_*iR*_= 4 × 10^−6^ M, and *V*_*mf*_= 0.05 mol s^-1^ (liter mito volume)^-1^. The parameter *V*_*mf*_is identified based on fitting model simulations to data.

#### Malic enzyme

Malic enzyme flux is simulated using an ordered bi-bi model:

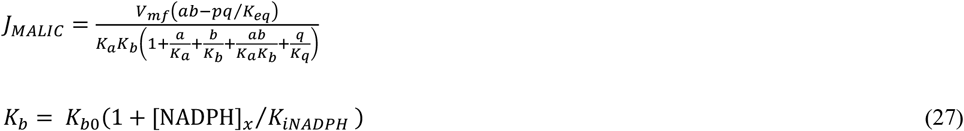

where *a* = [MAL]_x_, *b* =[NAD]_x_, *p* = [PYR]_x_, and *q* = [NADH]_free_. (Pyruvate terms in the denominator are assumed to be not significant.) It is assumed that NADPH competitively inhibits NAD binding. Parameter values are *K*_*a*_= 0.0877 × 10^−3^ M, *K*_*b*_= 0.63298 × 10^−3^ M, *K*_*q*_= 0.30388 × 10^−4^ M, *K*_*iNADPH*_= 0.51198 × 10^−3^ M, and *V*_*mf*_= 7.368 × 10^−5^ mol s^-1^ (liter mito volume)^-1^. All kinetic parameters are identified based on fitting model simulations to data.

#### Mitochondrial proton translocating transhydrogenase

The redox transfer from NAD(H) to NADP(H), which is electrogenically driven by inner membrane electrostatic potential. The overall reaction is

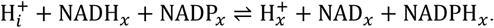

Flux is simulated using the following phenomenological expression

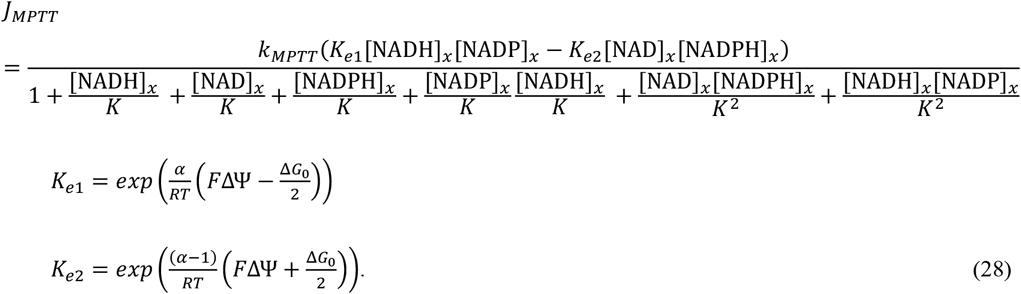

The constant *K* = 1 mM is arbitrarily set, and *k*_*MPTT*_= 1 mol s^-1^ (liter mito volume)^-1^ M^-2^.

#### Oxaloacetate decarboxylase

The model uses a simple Michaelis-Menten formulation for oxaloacetate decarboxylase flux:

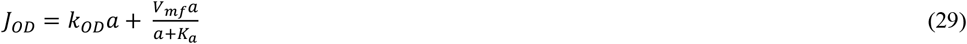

where *a* = [OAA]_x_and the first term represents spontaneous degradation of oxaloacetate. Parameter values *K*_*a*_= 14.52 × 10^−6^ M, and *V*_*mf*_= 4.3305 × 10^−5^ mol s^-1^ (liter mito volume)^-1^ are identified based on fitting model simulations to data. The first order rate constant *k*_*OD*_= 10 × 10^−4^ mol s^-1^ (liter mito volume)^-1^ M^-1^ is estimated based on observations of spontaneous degradation of oxaloacetate into pyruvate at 37ºC.

#### Phosphate-hydrogen co-transporter

A flux expression for the phosphate-hydrogen co-transporter is derived based on the random-order transporter model diagrammed in Figure A1.

The model has eight conformation and binding states, with configuration 1 representing conformational states with phosphate (p) and hydrogen ion (h) binding sites associated with the IM side of the mitochondrial inner membrane. The bindings site locations are flipped to the matrix side in configuration 2. The empty parentheses “()” indicate the location of the empty binding site in the two configurations.

Assuming rapid equilibrium binding we define

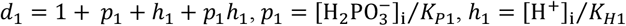

and

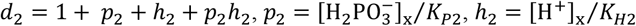

where *K*_*P*1_, *K*_*H*1_, *K*_*P*2_, and *K*_*H*2_are dissociation constants. Mass conservation requires

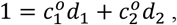

where 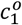 and 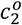 are the fractions of the transporter in configuration 1 and configuration 2.

Under steady-state conditions, the flux through the transporter may be expressed

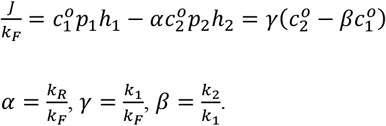

From these expressions we can solve for 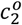 in terms of 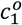:

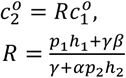

and thus

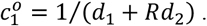

Finally, we get an expression for the transporter flux

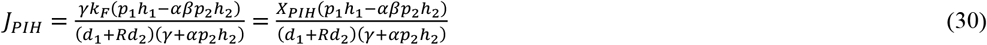

where *X*_*PIH*_represents the activity of the transporter. Parameters *γ* and *β* are assumed equal to 1. This implies a symmetry that *k*_1_= *k*_2_and that the rates of transition *k*_1_and *k*_*F*_are equal.

Thermodynamic reversibility requires

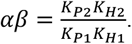

Parameter values are *K*_*P*1_= *K*_*P*2_= 1.8 × 10^−3^ M and *K*_*H*1_= *K*_*H*2_= 0.5 × 10^−7^M. The parameter *X*_*PIH*_= 0.35308 mol s^-1^ M^-2^ (liter mito volume)^-1^ is identified based on fitting model simulations to data.

#### Adenine nucleotide translocase

The adenine nucleotide translocase model is based on the model of Metelkin et al. [58], with parameter values updated by Bazil et al. [33]:

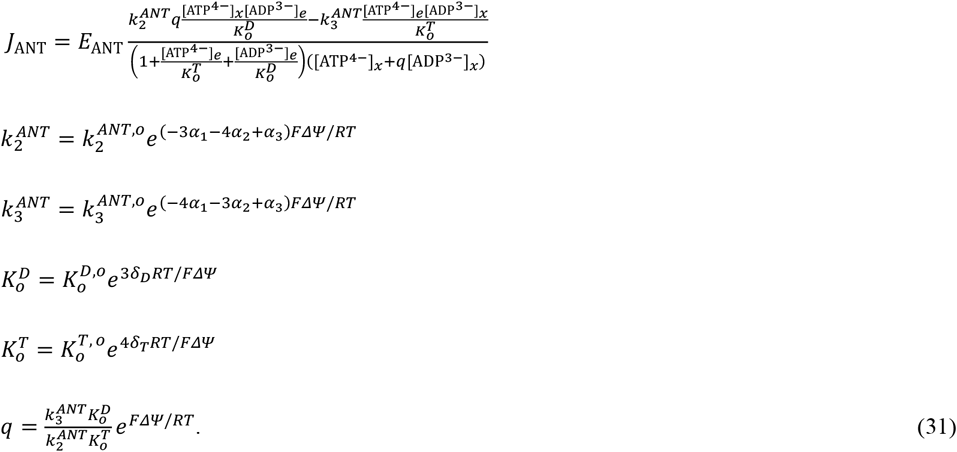

Parameter values are *E*_*ANT*_= 0.5398 mol (liter mito volume)^-1^, 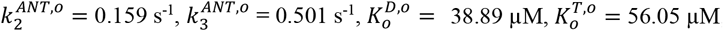, *α*_*1*_= 0.2829, *α*_*2*_= −0.2086, *α*_*3*_= 0.2372, *δ*_*T*_= 0.0167, and *δ*_*D*_= 0.0699. The parameter *E*_*ANT*_is identified based on fitting model simulations to data.

#### H^+^ and K^+^ leaks

The hydrogen and potassium ion passive leak currents are assumed to be governed by the Goldman-Hodgkin-Katz

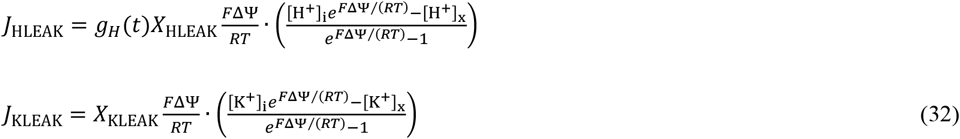

where *X*_HLEAK_= 4872.9 mol s^-1^ (l mito)^-1^ M^-1^ and *X*_KLEAK_= 1.2316 × 10^−3^ mol s^-1^ (l mito)^-1^ M^-1^ are the maximal conductivities, and are treated as adjustable parameters. The potassium ion leak is assumed constant while the hydrogen ion conductivity varies according to Equation (7).

#### Mitochondrial K^+^ / H^+^ exchanger

The hydrogen-potassium exchange flux is modeled using the simple mass-action relationship

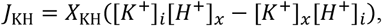

where the activity *X*_KH_= 1.2458 × 10^6^ mol s^-1^ (l mito)^-1^ M^-2^ is treated as an adjustable parameter.

#### Pyruvate-H^+^ co-transporter

Pyruvate transporter is governed by the monocarboxylate transporter model of Vinnakota and Beard [59]:

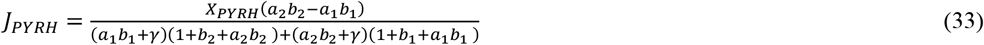

where *a*_1_= [PYR]_x_⁄*P*_*PYR,x*_*K*_*PYR*_, *h*_1_= [H^+^]_x_⁄*K*_*H*_, *a*_2_= [PYR]_i_⁄*P*_*YR,i*_*K*_*PYR*_, and *h*_2_= [H^+^]_i_⁄*K*_*H*_. We assume the value of *γ* = 2.3 based on the value estimated for the lactate transporter [59]. Moreover, the kinetic parameters *K*_PYR_= 0.071631 × 10^−3^ M and *K*_H_= 10^−7^ M are set to arbitrary values. The overall transporter activity *X*_PYRH_= 0.052365 mol s^-1^ (l mito)^-1^ M^-2^ is treated as an adjustable parameter and identified based on fitting model simulations to data.

#### Glutamate-H^+^ co-transporter

A simple mass-action model is used to govern glutamate-H^+^ co-transport:

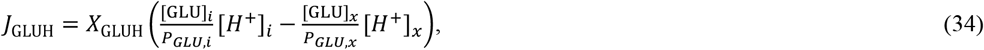

where overall transporter activity *X*_GLUH_= 1.9213 × 10^6^ mol s^-1^ (l mito)^-1^ M^-2^ is treated as an adjustable parameter and identified based on fitting model simulations to data.

#### Citrate/malate antiporter

Citrate-malate antiport is not accounted for in the current model simulations:

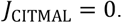

#### Dicarboxylate transporter: succinate/phosphate and malate/phosphate antiport

A flux expression for the succinate-phosphate and malate-phosphate exchange through the dicarboxylate transporter is derived based on alternating-access mechanism diagrammed in Figure A2.

The model has 8 conformational states, with configuration 1 representing conformational states with the binding site associated with the IM side of the mitochondrial inner membrane. The dicarboxylate/phosphate binding site location is flipped to the matrix side in configuration 2. The empty parentheses “()” indicate the location of the empty binding site in the two configurations.

Assuming rapid equilibrium binding we define

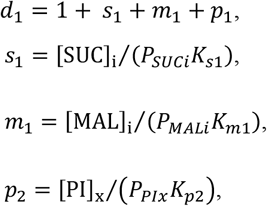

where *P*_*SUCi*_, *P*_*MALi*_, *P*_*PIx*_are the binding polynomials for IM succinate, IM malate, and matrix phosphate, and *K*_*s*1_, *K*_*m*1_, and *K*_*p*2_are dissociation constants associated with configuration 1. Similarly, we define

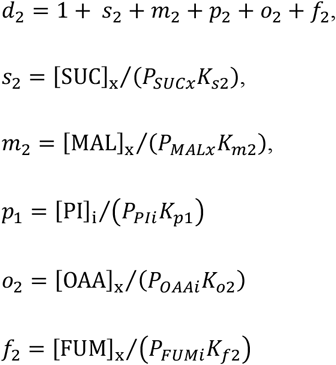

where *P*_*SUCx*_, *P*_*MALx*_, *P*_*PIx*_are the binding polynomials for matrix succinate, matrix malate, and IM phosphate, and *K*_*s*2_, *K*_*m*2_, and *K*_*p*1_are dissociation constants associated with configuration 2. The *o*_2_and *f*_2_terms are associated with inhibitory binding of the transporter with matrix oxaloacetate and fumarate.

Defining 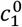 and 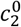 as the relative proportion of transporter in unbound state in configurations 1 and 2, we have

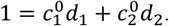

With these definitions we can compute the relative proportion of transporter in any of its states. For example, the proportion of transporter in configuration 1 with phosphate bound is computed 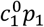. Steady-state flux balance requires

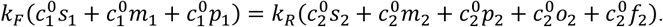

Solving for 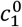 and 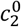, we obtain

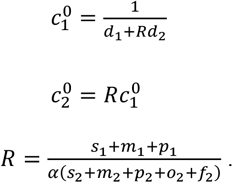

where *α* = *k*_*R*_⁄*k*_*F*_.

The net transport of succinate from IM to matrix space is computed

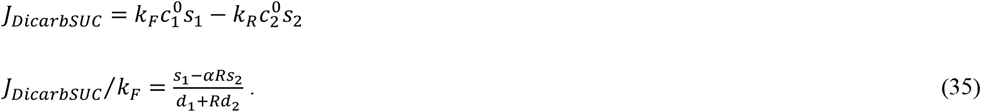

Similarly,

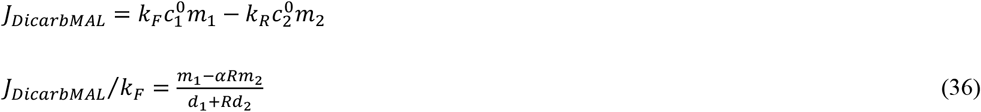

and

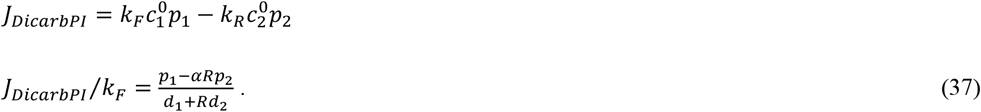

Parameter values *k*_*F*_= 9.0417 × 10^−3^mol s^-1^ M^-2^ (liter mito volume)^-1^, *K*_*S*1_= 3.114 × 10^−3^ M, *K*_*m*1_= 4.101 × 10^−3^ M, *K*_*p*1_= 6.240 × 10^−3^ M, *K*_*o*2_= 1.267 × 10^−6^ M, *K*_*f*2_= 1.303 × 10^−3^ M are estimated by fitting model simulations to data. (Parameter and *α* is set to 1.)

Parameters *K*_*p*2_, *K*_*s*2_, *K*_*m*2_, and *K*_*o*2_are computed from thermodynamic reversibility constraints

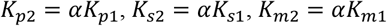

#### α-Ketoglutarate/malate antiporter

The oxoglutarate carrier, facilitating α-ketoglutarate/malate antiport, follows an alternating-access mechanism diagrammed in Figure A3.

The model has 6 conformational states, with configuration 1 representing conformational states with the binding site associated with the IM side of the mitochondrial inner membrane. The binding site location is flipped to the matrix side in configuration 2. The empty parentheses “()” indicate the location of the empty binding site in the two configurations.

Assuming quasi-equilibrium binding, we introduce

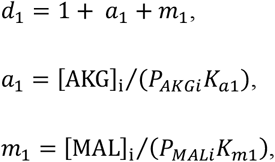

where *P*_*AKGi*_and *P*_*MALi*_are the binding polynomials for IM α-ketoglutarate and malate, and *K*_*a*1_and *K*_*m*1_are dissociation constants. Similarly, we define

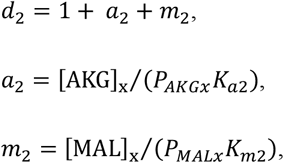

where *P*_*AKGx*_and *P*_*MALx*_are the binding polynomials for matrix α-ketoglutarate and malate, and *K*_*a*2_and *K*_*m*2_are dissociation constants.

Similar to the formulation for the dicarboxylate transporter, we have

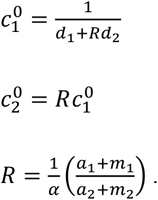

where *α* = *k*_*R*_⁄*k*_*F*_.

The net transport of malate and α-ketoglutarate from the IM to matrix space is computed

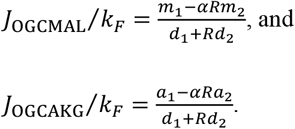

Furthermore, we have

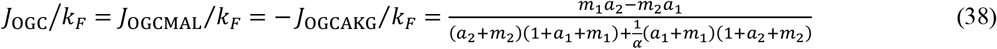

Parameter values are *k*_*F*_= 0.01228 mol s^-1^ (liter mito volume)^-1^, *K*_*a*1_= 1.0 × 10^−3^ M, *K*_*m*1_= 1.0 × 10^−3^ M, and *α* = 1. (Parameters *K*_*a*1_, *K*_*m*1_, and *α* are set to arbitrary values and not treated as adjustable parameters.)

Parameters *K*_*a*2_, and *K*_*m*2_are computed from thermodynamic reversibility constraints

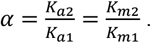

#### Aspartate/glutamate carrier

The aspartate/glutamate carrier, facilitating aspartate/glutamate antiport, follows an alternating-access mechanism similar to the dicarboxylate transporter and the oxoglutarate carrier. A key difference is that the aspartate/glutamate exchange is electrogenically driven by the inner membrane electrostatic potential. To develop a model for the transporter, consider the mechanism illustrated in Figure A4:

Following an analogous derivation, we define

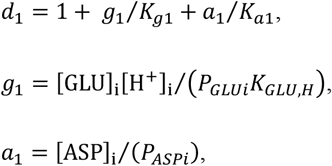

where *P*_*GLUi*_and *P*_*AKGi*_are the binding polynomials for IM glutamate and aspartate, and *K*_*g*1_and *K*_*a*1_are dissociation constants. Similarly,

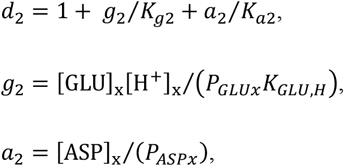

where *P*_*GLUi*_and *P*_*AKGi*_are the binding polynomials for matrix glutamate and aspartate, and *K*_*g*2_and *K*_*a*2_are dissociation constants.

As for other transporters above, we define 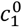 and 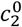 as the relative proportion of transporter in unbound state in configurations 1 and 2:

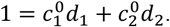

With these definitions we can compute the relative proportion of transporter in any of the 8 states. For example, the proportion of transporter in configuration 1 with phosphate bound is computed 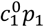. Steady-state flux balance requires

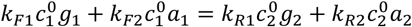

Solving for 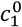 and 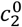,

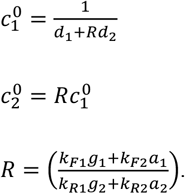

With the following constraints

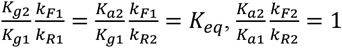

the net turnover of the aspartate glutamate carrier, with glutamate transported from IM to matrix space, simplifies to

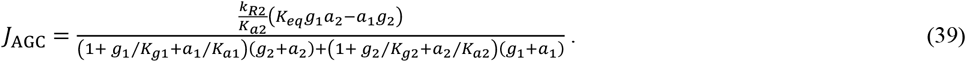

Since this transport mechanism is associated with the net transfer of one charge from the IM to matrix space, the equilibrium constant is computed

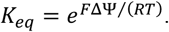

Parameter values are *k*_*R*2_⁄*K*_*a*2_= 91.744 × 10^−6^ mol s^-1^ (liter mito volume)^-1^, *K*_*g*1_= 0.96899 × 10^−3^ M, *K*_*a*1_= 0.70257 × 10^−3^ M, *K*_*g*2_= 0.77793 × 10^−3^ M.

#### Ammonia flux across inner membrane

Ammonia is assumed to be passively transported out of the matrix

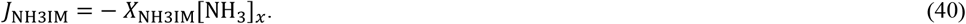

The activity is set to *X*_NH3IM_= 1 mol s^-1^ (liter mito volume)^-1^ M^-1^. This value is arbitrary because matrix reactions are assumed to not be influenced by ammonia concentration.

#### Outer membrane permeability

Passive permeation across the outer membrane is governed by

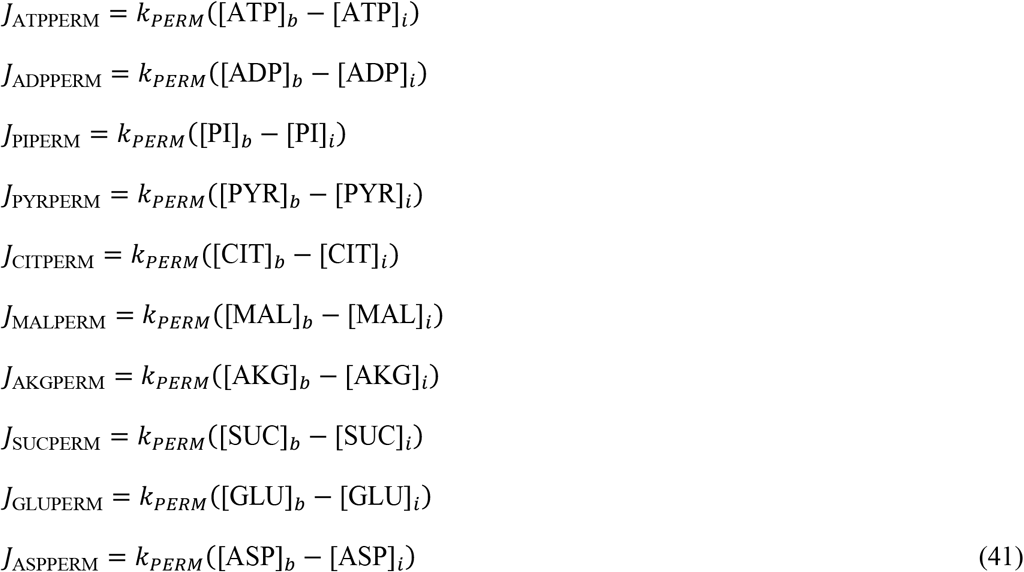

The value *k*_*PERM*_= 400 mol s^-1^ (liter mito volume)^-1^ M^-1^ ensure rapid equilibration of concentrations across the outer membrane.

#### Buffer ATP hydrolysis

The ATPase rate expression is adopted from Bazil et al. [33]:

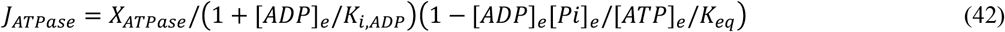

where *K*_*i,ADP*_= 0.10 × 10^−3^ M. The ATP hydrolysis activity is estimated to be *X*_*ATPase*_= 0.38 × 10^−6^ mol s^-1^ (liter mito volume)^-1^.

#### Buffer adenylate kinase reaction

A mass-action expression is used to maintain adenylate kinase equilibrium mass-action ratio:

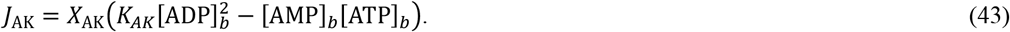

The activity *X*_*AK*_= 2 mol s^-1^ (liter mito volume)^-1^ M^-2^ is arbitrarily set at a value high enough to effectively maintain adenylate kinase equilibrium.

---

**Figure A1:**
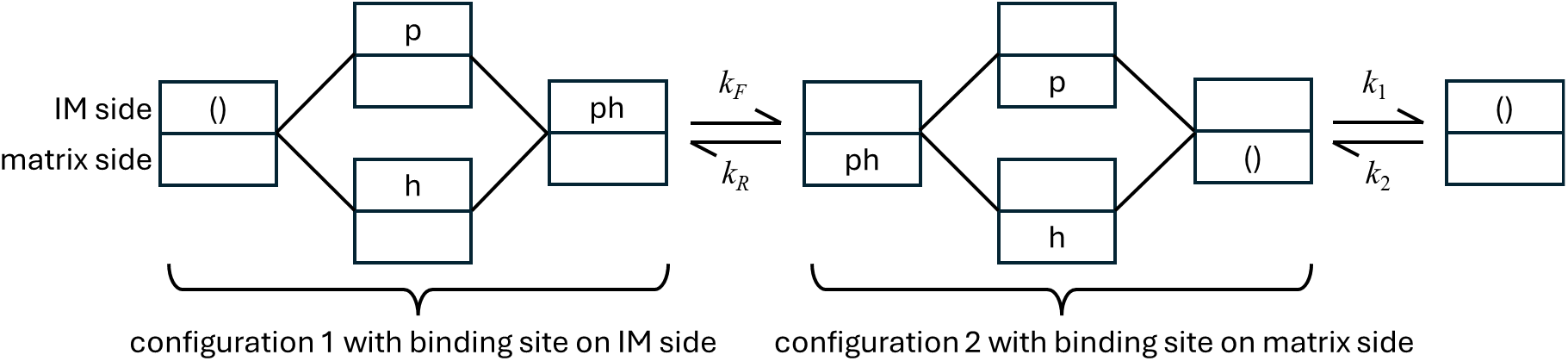
Diagram of phosphate-hydrogen co-transporter model.

**Figure A2:**
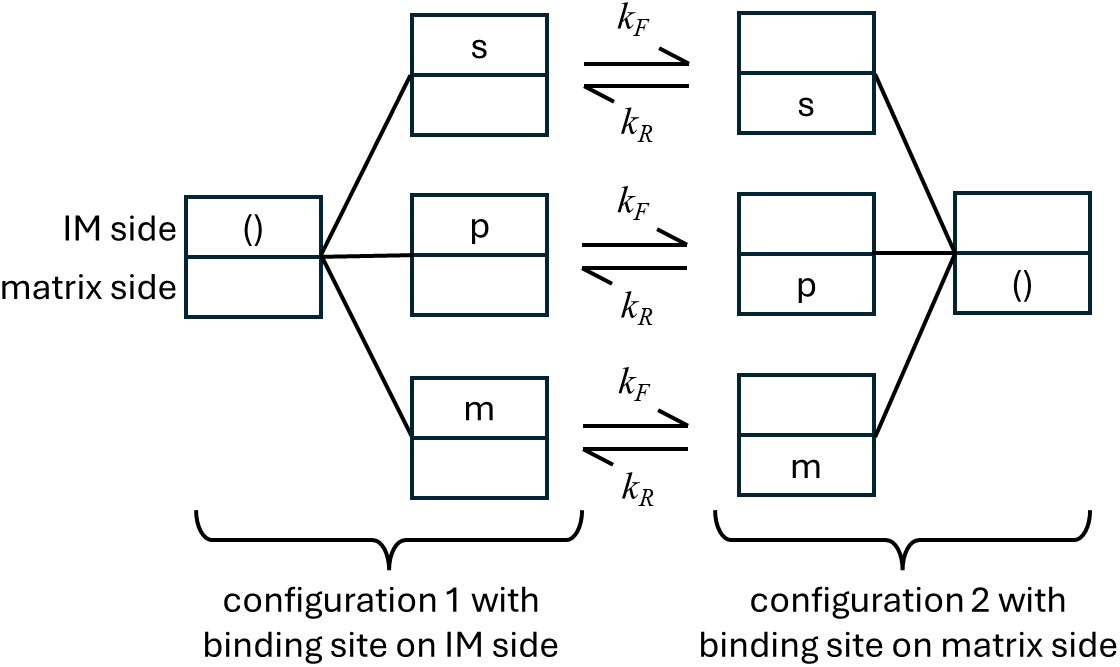
Diagram of dicarboxylate transporter model.

**Figure A3:**
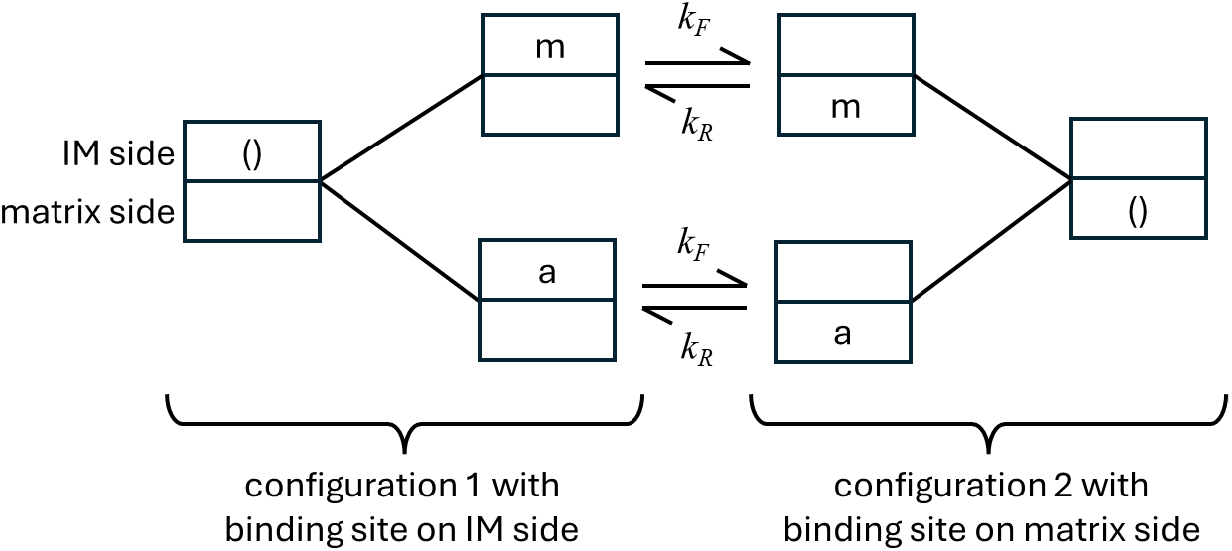
Diagram of oxoglutarate carrier model.

**Figure A4:**
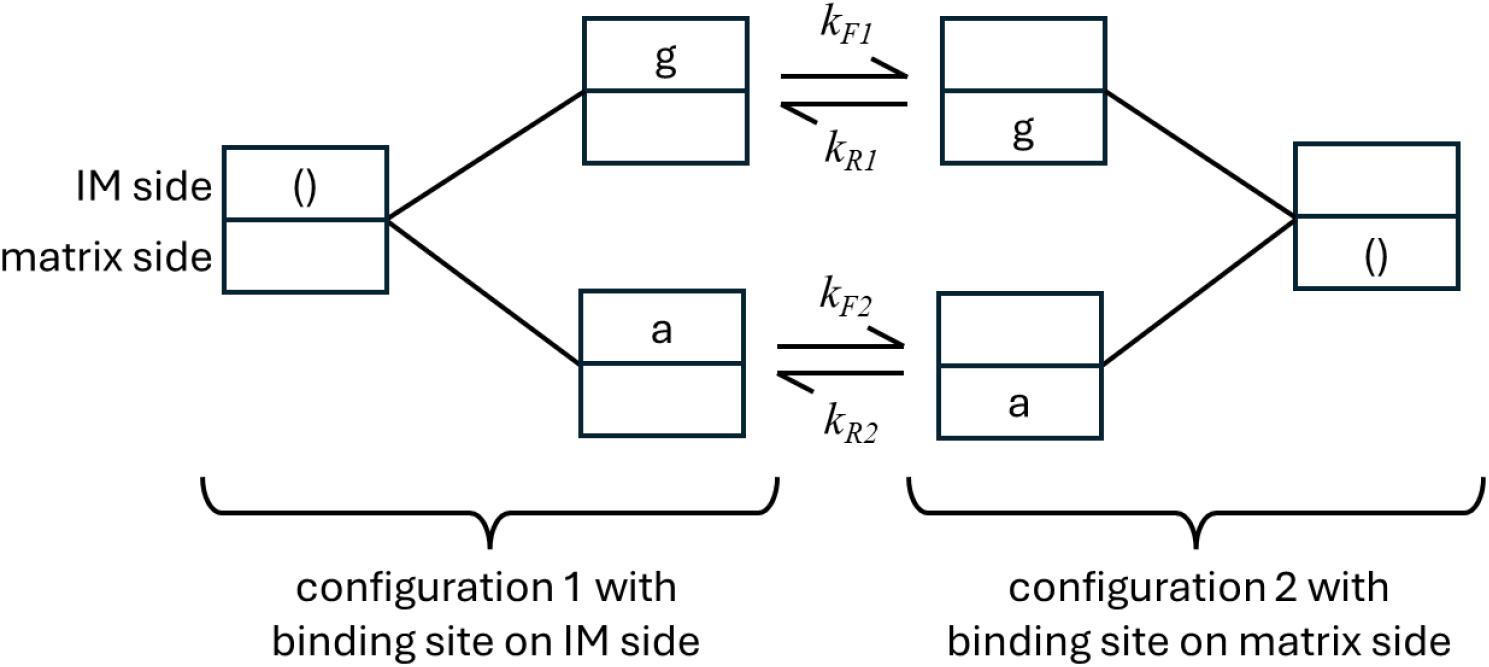
Diagram of aspartate-glutamate carrier model.

